# Benchmarking fragmentation-derived artificial cfDNA reference standards

**DOI:** 10.64898/2026.08.12.744389

**Authors:** Lotte Cornelli, Truong Nhat Nguyen, Robin Van Belle, Sofie Roelandt, Aaron De Cock, Joni Van der Meulen, Siebe Loontiens, Nadine Van Roy, Katleen De Preter

## Abstract

An important step toward clinical implementation of (epi-)genomic assays on liquid biopsies is their validation on identical samples within and across laboratories. For these validation studies, there is a need for cell-free DNA (cfDNA) samples with defined tumor fractions and (epi-)genomic aberrations. However, the amount of circulating cfDNA isolated from patient samples is often limited, especially in pediatric cases. Additionally, patient samples contain a high degree of variability in cfDNA yield and tumor fraction. Several commercial artificial cfDNA products are available for validation studies, however their use is restricted to specific assays, aberrations and/or tumor entities. Alternatively, artificial cfDNA samples can be produced by fragmenting genomic DNA to mimic highly fragmented cfDNA derived from both tumor and healthy blood, followed by mixing artificial tumoral and healthy cfDNA at defined fractions. In this study, we compared native cfDNA with artificial cfDNA generated by three different fragmentation methods, including sonication and two enzymatic digestions using micrococcal nuclease and double-stranded deoxyribonuclease (dsDNase). We assessed fragment length profiles, end motifs and nucleosome occupancy patterns from shallow whole-genome sequencing data, as well as coverage profiles from targeted panel sequencing, together with a small-scale mixing experiment of tumor and healthy cell derived artificial cfDNA. Although sonication remains a convenient high-throughput approach to generate artificial cfDNA for certain downstream applications, enzymatic fragmentation, particularly the dsDNase-based method, more faithfully reproduced native cfDNA characteristics.

## Introduction

Molecular profiling of tumors is an important cornerstone in the routine diagnostic process, supporting accurate diagnosis, prognosis and personalized treatment decisions^1–3^. However, these assays require invasive tissue biopsies that only inform about the sampled region, posing challenges for heterogeneous tumors and limiting repeated sampling during follow-up. In addition, biopsies may yield only limited amounts of tissue, which can be largely or completely exhausted during routine histopathological evaluation, leaving insufficient material for comprehensive molecular testing. As a result of cell death processes, mostly apoptosis, tumor biomolecules are shed into surrounding biofluids such as blood, urine, and saliva^4–6^. Collecting these biofluids, or liquid biopsy samples, gives direct access to cell-free DNA (cfDNA), extracellular RNA, circulating proteins and extracellular vesicles released by tumor cells^7–9^. Therefore, molecular profiling of liquid biopsy samples of cancer patients offers a minimally-invasive alternative to tissue-based profiling^10–12^. Several plasma cfDNA methods are implemented in clinical practice for DNA mutation detection and copy number profiling^13–17^,and new fragmentomic and epigenetic approaches are under development^18–22^.

Circulating molecules are also released by healthy cells, therefore cfDNA in cancer patients is a mixture of tumor-derived and healthy DNA. When analyzing specific genomic aberrations, healthy cell derived cfDNA can dilute the tumoral signal, therefore false negative tests can occur, and more sensitive variant detection methods are necessary. Assay performance depends on the tumoral cfDNA fraction, with lower tumoral cfDNA fractions compromising sensitivity. Importantly, the tumoral cfDNA fraction is also an informative biomarker associated with disease recurrence, tumor burden, and aggressiveness^23,24^. To validate cfDNA assays before clinical implementation, reference samples with defined aberrations and tumor fractions are needed. Using samples with varying tumor fractions allows determining the lowest fraction at which aberrations can still be detected. Several interlaboratory comparison studies, also termed *Round Robin Studies* or *Ring Trials,* have used different strategies to obtain reference samples^25–29^. Some used patient samples with high volume and known genomic aberrations^26,27^, e.g. by aliquoting large plasma volumes, or by collecting plasma from multiple patients and randomizing each patient sample in at least two different centers. However, these approaches are less feasible for cases with low plasma availability such as pediatric cancer patients and are statistically complex when multiple centers are involved. Additionally, patient cfDNA samples show high variability in tumor fraction across tumor types, patients, and time points^30–34^. Moreover, subclonal tumor populations may lack certain aberrations, resulting in heterogeneous cfDNA mixtures in which the fraction of molecules carrying a given aberration differs from the overall tumor fraction^35,36^. Together, these factors complicate the use of patient samples for assay validation.

Alternatively, artificial cfDNA samples with defined tumor fraction, i.e. controlled mixtures of healthy and tumoral DNA, can be used to validate cfDNA assay performance. Ideally, cfDNA reference samples should resemble native cfDNA fragmentation, typically around 160 bp fragments (or multiples) due to apoptosis^37,38^. More specifically, cfDNA is fragmented by deoxyribonuclease (DNase) enzymes that cannot digest the DNA wrapped around histones, producing a pattern with mono-nucleosomal peaks and less abundant di-or tri-nucleosomal peaks. Cell-fee DNA reference samples can be either commercially available^28,39^, or generated in house by sonication^25,29^ or enzymatic fragmentation^40–44^. During sonication, sound waves mechanically fragment DNA strands into lengths corresponding to cfDNA. Sonication results in random fragmentation and therefore fundamentally differs from the cfDNA fragments generated by cell death processing including apoptosis^4,45^. Alternatively, enzymatic fragmentation using micrococcal nuclease (MNase) or double-stranded deoxyribonuclease (dsDNase) results in fragments that better resemble nucleosomal cfDNA^46,47^. Applying these approaches on white blood cells from healthy donors and tumor cells from cancer cell lines allows generation of cfDNA mixtures with known tumor fraction.

In this study, we aimed to provide a comprehensive overview of artificial cfDNA models and investigate how well they recapitulate circulating native cfDNA in human plasma. We first evaluated commercially available reference cfDNA samples for cancer diagnostic assay validation. In addition, we assessed cfDNA isolated from plasma alongside artificial cfDNA generated by MNase, dsDNase fragmentation and sonication. Using shallow whole-genome sequencing (sWGS) data, we evaluated key molecular features including fragment length profiles, end motifs and nucleosome occupancy. In addition, we performed panel sequencing to assess coverage variability across the included regions. A small-scale mixing experiment with healthy and tumor cell derived artificial cfDNA was performed to validate the results in a mixed context.

## Results

### Commercial reference material for cfDNA assay validation studies

Before comparing in-house methods for artificial cfDNA generation, artificial cfDNA references available from commercial suppliers were listed and reviewed. These DNA references are summarized in Table 1. Half of the listed products are generated by spiking synthetic DNA into a healthy cfDNA background, confining their application to detecting the spiked genomic aberrations (SNVs, INDELs, CNVs, SVs). The healthy background is obtained through fragmentation of wild-type genomic DNA or DNA from non-cancerous cell lines. In most cases transparency is lacking in the fragmentation details, and some use patented approaches. While most suppliers previously provided sonicated reference material, many have switched or plan to switch enzymatic methods enabling the validation of epigenomic-based cfDNA analysis including nucleosomics. Most suppliers provide reference material for common genomic aberrations and tumor types, resulting in underrepresentation of rare cancers (e.g. pediatric cancers) or rare aberrations and thereby necessitating in-house protocols for artificial cfDNA generation using cells of interest with known (epi)genomic aberrations.

**Table 1.** Overview of commercial suppliers of cfDNA reference or cfDNA in plasma reference, including information on the cfDNA preparation, healthy background, presence of genomic aberrations, tumor type applicability and potential to investigate epigenomic information (VAF = Variant Allele Fraction, SNV = single nucleotide variant, CNV = Copy number variation, INDEL = Insertion and Deletion, SV = Structural Variant).

| Supplier | Product | cfDNA reference preparation and genomic aberrations | Fragment length & fragmentation approach | Tumor type | Epigenomic analysis potential, e.g. methylation/histone modifications |
| --- | --- | --- | --- | --- | --- |
| **Sera Care (LGC Clinical diagnostics)**<sup>54</sup> | *Seraseq® ctDNA mutation mix* | Synthetic DNA with known mutations diluted into a background of wild-type genomic DNA, at different VAFs: 40 clinically-relevant mutations across 28 genes. Other reference material from Sera Care contains INDELs, CNVs, SVs. | ~170 bp (synthetic DNA and patented method of fragmentation) | Not tumor specific. | No epigenomic information of the tumor. Sera Care also distributes reference material with all or none CpG's methylated. |
| **Horizon Discovery** | *Mimix OncoSpan cfDNA Reference Standard (DNA reference)* | Cell line derived reference material with known mutations at different VAFs: over 380 variants across 152 oncogenes. | ~160 bp (sonication, but investigating enzymatic approaches) | Not tumor specific. Horizon Discovery also distributes tumor specific reference material. | Methylation retained but other epigenomic information lost with the sonicated material. |
| **Twist Bioscience**<sup>55</sup> | *Twist pan-cancer cfDNA Reference Standard (DNA reference)* | Wild-type (WT) background cfDNA from a cell-line and synthetic oligos carrying mutant alleles at different VAFs: 400 variants across 84 genes (incl. SNVs, INDELs, SVs). | ~167 bp (synthetic DNA and cell line derived cfDNA: not specified) | Not tumor specific. | No epigenomic information of the tumor. |
| **Thermo Fisher/ ArcoMetrix**<sup>56</sup> | *AcroMetrix™ Multi-Analyte ctDNA Plasma Control (DNA in plasma reference)* | Fragmented synthetic DNA and genomic DNA from GM24385 human cell line in normal human plasma matrix at different VAFs. 13 variants including SNVs, INDELs, and CNVs. | cfDNA-like fragments, not further specified | Broad range of cancer types: breast, colon, lung, ovaries, pancreas, skin, thyroid, and more | No epigenomic information of the tumor. |
| **nRich**<sup>DX 44</sup> | *cfDNA Reference Standard (DNA reference to be spiked in plasma)* | Enzymatic fragmentation using a nucleosome preparation kit on NCI-H441 lung cancer cell line: KRAS G12V and TP53 Arg158Leu | ~150bp (enzymatic fragmentation of NCI-H441 lung cancer cell line) | Lung cancer | Epigenomic information of the tumor retained. |
| **ZeptoMetrix**<sup>57</sup> | *cfDNA controls (DNA reference as well as DNA in plasma reference)* | Mutated synthetic DNA, mixed in healthy isolated cfDNA: 50 clinically relevant somatic mutations at 3% VAF, pre-spiked into DNA-depleted human plasma, or provided as DNA. | Not disclosed | Not tumor specific. Reference for ESR1 mutations | Not disclosed |
| **Anchor Molecular**<sup>58</sup> | *ctDNA standard (DNA in plasma reference)* | 23 multiplexed nucleosomal fragmented ctDNA fragments (23 cell derived variants) mixed with normal patient cfDNA background in human plasma. | ~150 bp (generated by Anchor's unique multiplexed gene-editing method and are nucleosomal fragmented) | Not tumor specific | Not disclosed |
| **SensID<sup>59</sup>** | *5-gene*<br>*multiplex*<br>*cfDNA*<br>*Nucleosomal*<br>*healthy DNA*<br>*EGFR/5-Gene*<br>*multiplex*<br>*cfDNA* | Healthy DNA fragmented<br>nucleosomal, mixed with<br>enzymatically fragmented DNA from<br>lung cancer cell line containing<br>mutations<br>AKT1/BRAF/ERBB2/KRAS/PIK3CA<br>*DNA reference to be spiked in plasma* | ~167bp | Specific mutations | Histones and DNA<br>methylation potentially<br>conserved. |

### Artificial cfDNA generation and fragment size analysis

Blood was collected from four healthy donors, cfDNA was isolated from plasma and the PBMC-DNA was fragmented using mechanical fragmentation (i.e. sonication) and enzymatic fragmentation (using dsDNase or MNase). Fragment size profiles (Tapestation) of one representative sample per condition is shown in Figure 1. Enzymatically digested DNA samples show a two-peak pattern comparable to mono-and dinucleosomal peaks observed in plasma cfDNA, though the dinucleosomal peak is more subtle. The length of the mononucleosomal peak is longer after dsDNase digestion than after MNase digestion. MNase-treated samples show smaller fragments, that are not observed in the dsDNase-treated samples. As expected, the sonicated samples with a random fragmentation pattern show a smooth distribution with a 188 bp peak prior to library preparation.

**Figure 1:**
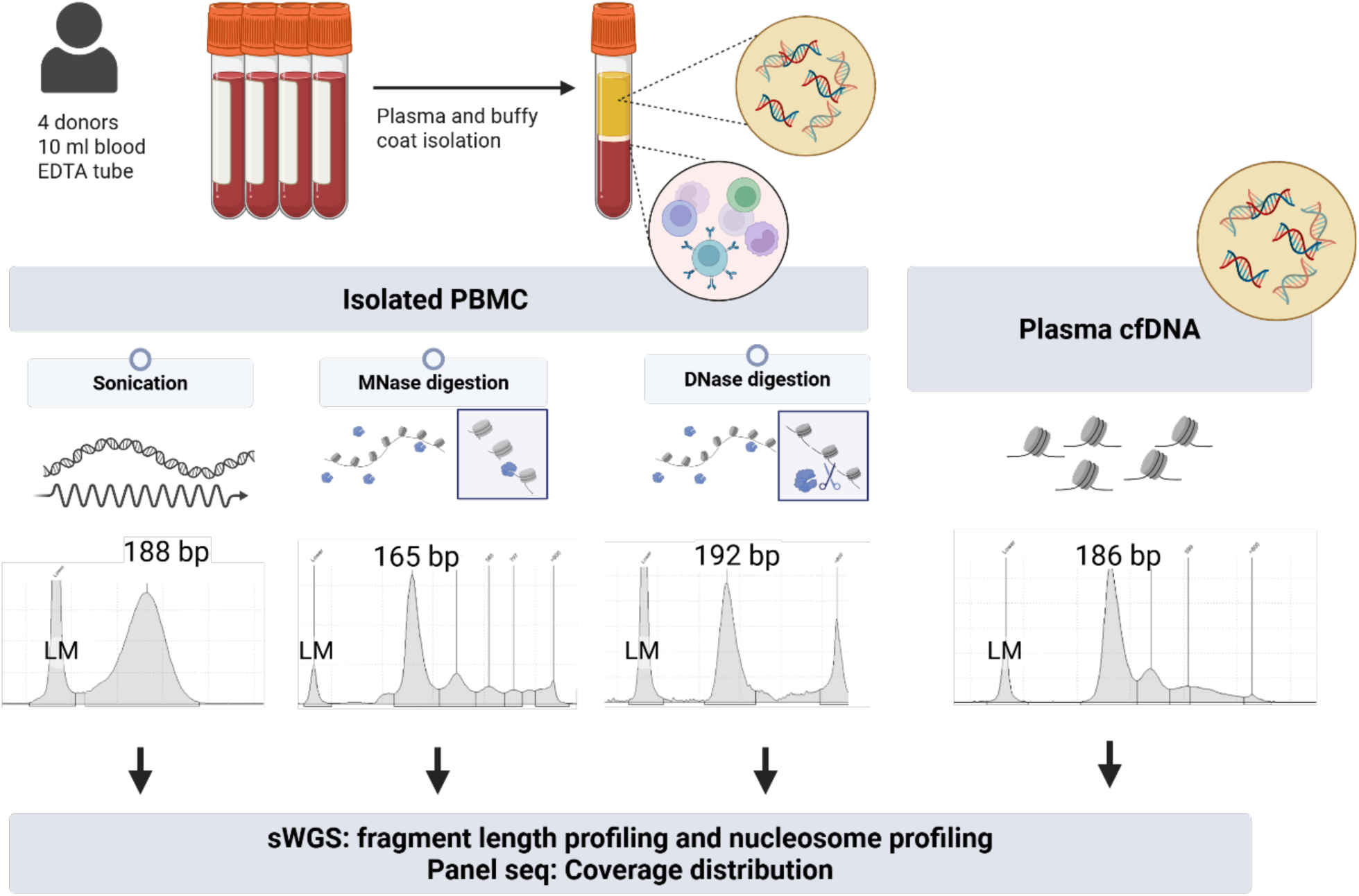
Experimental overview. Blood was collected from 4 healthy donors. Plasma was separated from the buffy coat. cfDNA was isolated from plasma samples. From the buffy coat, PBMCs were isolated. The DNA present in live PBMCs was enzymatically fragmented by MNase and dsDNase. In parallel, DNA was isolated from the PBMCs and sonicated to a measured length of 188bp, corresponding to the measured length of cell-free DNA of 186bp (assuming a fragment length overestimation by the TapeStation). sWGS and panel sequencing was performed on all the samples to investigate the fragmentomic patterns, nucleosome profile and the coverage distribution over the gene panel. PBMC: Peripheral Blood Mononuclear Cells; MNase: micrococcal nuclease; sWGS: shallow Whole-Genome Sequencing; Panel seq: panel sequencing, LM: lower marker. The figure was created with *BioRender.com*

### Artificial fragmentation methods generate distinct fragment length and end-motif profiles compared to plasma cfDNA

### Fragment length profiles

Shallow WGS was performed on artificial cfDNA and plasma cfDNA to compare their fragmentomic features, including fragment length distributions and the ratio of short fragments (100-150 bp) to mononucleosome-sized fragments (150-200 bp). Plasma cfDNA showed a dominant ∼166 bp peak and a minor ∼330 bp peak, consistent with mono-and dinucleosomal fragments, together with ∼10 bp periodicity among short fragments in the 50–150 bp range, supporting nucleosome-associated fragmentation (Figure 2a). In contrast, enzymatic and mechanical fragmentation of PBMC DNA produced distinct profiles. dsDNase digestion generated a dominant ∼140 bp peak and multiple short fragment peaks between 60 and 160 bp with ∼10 bp periodicity. MNase digestion enriched shorter fragments (50–90 bp) together with a prominent ∼140 bp peak, consistent with endonuclease or exonuclease digestion of chromatin associated DNA^53^. The shorter insert sizes observed in the dsDNase-digested libraries may be explained by the blunt ends generated during enzymatic digestion, which can result in shorter DNA fragments being preferentially retained during library preparation. Sonication produced a broader fragment-size distribution centered around ∼100–120 bp, reflecting more random fragmentation and, consequently, a loss of the characteristic chromatin-associated fragmentation patterns. The shorter insert sizes observed in these libraries may be related to single-stranded nicks introduced by sonication, which can prevent longer fragments containing such nicks from being efficiently amplified during PCR.

**Figure 2.**
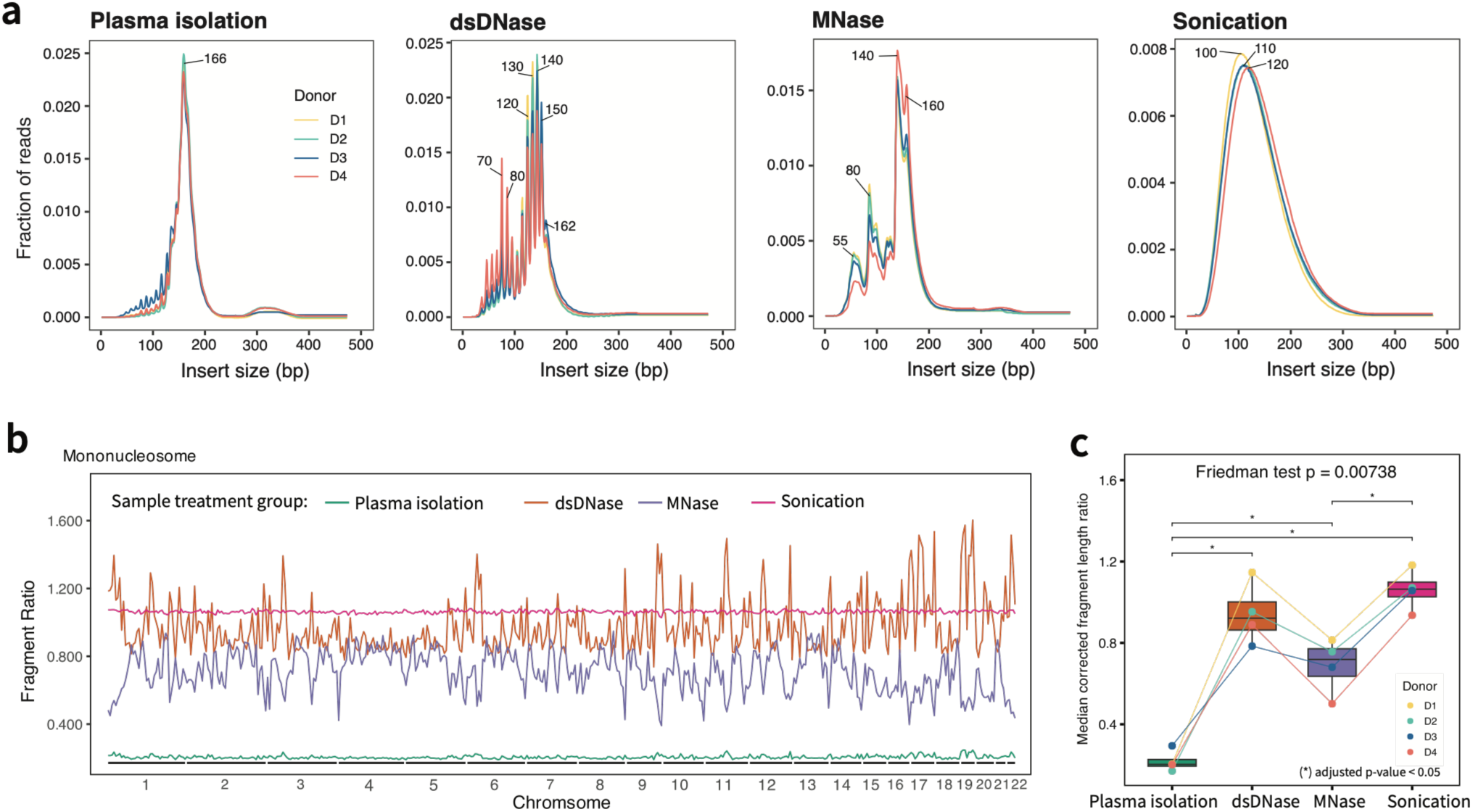
Fragment length profiles across donor-matched sample preparation methods. (a) Insert-size distributions for plasma isolation, dsDNase, MNase, and sonication samples. Lines indicate individual donors (D1–D4). (b) Genome-wide mononucleosomal fragment ratio across chromosomes for each treatment method. Fragment length ratio was calculated as the number of short fragments (100-150 bp) divided by the number of mononucleosome-sized fragments (151-200 bp) within each genomic bin. (c) Median corrected fragment length ratio compared across donor-matched treatment groups. Points represent individual donors, and lines connect matched donors.

Genome-wide short-to-mononucleosome fragment ratio profiles differed across conditions (Friedman test, p = 0.007) (Figure 2b, c). Plasma-derived cfDNA showed lower ratios compared to dsDNase, MNase, and sonication (all BH-adjusted p = 0.043). Together, these data indicate that artificial cfDNA samples exhibit a size profile that is similar, but not identical to plasma cfDNA, including mono-and dinucleosomal peaks when enzymatic fragmentation is used, but with fragment ratio profiles that are significantly different.

### Fragment end-motif profiles

As end-motif analysis of cfDNA fragments can reveal the tissue of origin, the type of cell death involved, and disease-associated signatures, motifs of the synthetic cfDNA samples were compared with those of plasma derived cfDNA. Motif profiling at 5’ end revealed marked differences in 1, 2 and 4-mer sequence composition across four sample treatment conditions (Figure 3a and Supplementary Figure 1). Plasma-isolated cfDNA displayed a heterogeneous end-motif landscape, with enrichment of motifs starting with cytosine (C). dsDNase-treated samples also showed predominance of C-starting motifs; however the enriched sequences differed from plasma cfDNA and showed a low proportion of thymine (T) starting motifs. MNase digestion generated a distinct 5′ motif profile, with prominent enrichment of adenine (A)-starting motifs, and abundant T-starting motifs, while C and G starting motifs were depleted. Sonication produced a broader and less biased motif distribution. Hierarchical clustering of 5’ end-motif z-scores separated samples primarily by treatment condition rather than donor group, indicating that plasma specific end motifs are not recapitulated in the artificial cfDNA samples (Figure 3b).

**Figure 3.**
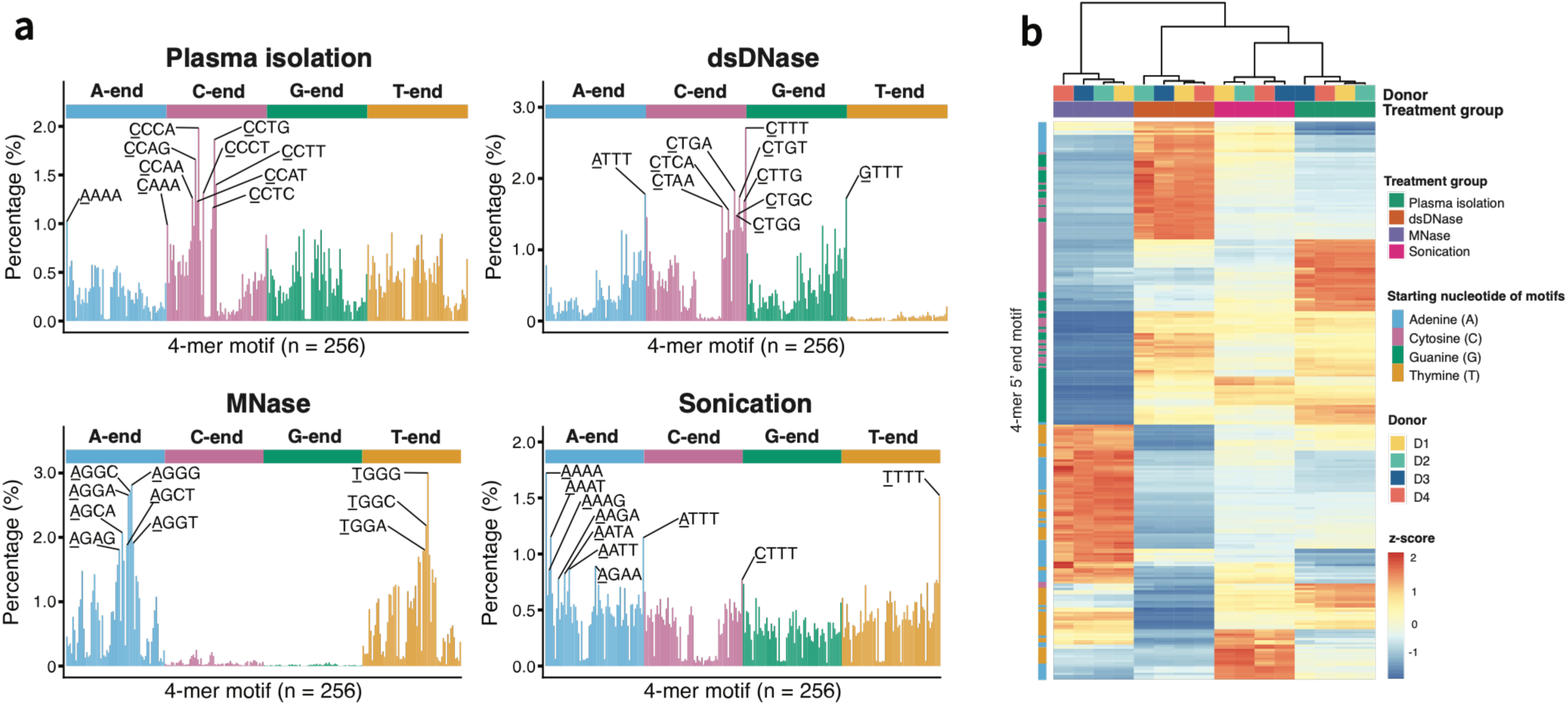
Fragment end-motif composition differs across plasma derived cfDNA and artificial cfDNA. (a) The fraction of 256 end motifs, ordered in alphabetical order for the 4 different sample types. The top 10 4-mer motifs with highest frequencies are highlighted and written in a 5’-to-3’ direction, with the base nearest to the cleavage site underlined. (b) Unsupervised hierarchical clustering of samples based on the frequencies of all 256 end motifs. Rows represent 4-mer 5’ end motifs and columns represent samples; values are shown as row-scaled z-scores.

### dsDNase fragmentation partially preserves cfDNA nucleosome signatures

To evaluate whether PBMC derived DNA fragmentation preserves cfDNA nucleosome features, we compared coverage profiles around binding sites of 270 transcription factors (TFs) across plasma cfDNA and artificial cfDNA using Griffin on sWGS data. Figure 4a shows coverage profiles centered on CTCF sites after GC correction, revealing the characteristic oscillatory pattern of plasma cfDNA consistent with phased nucleosome positioning^50^. A comparable periodic pattern appeared after dsDNase and MNase treatment. dsDNase-treated DNA retained clear oscillatory patterns of nucleosome, similar to plasma cfDNA, whereas MNase-treated DNA showed pronounced central enrichment and sharper periodic peaks. In contrast, sonicated DNA displayed a disrupted profile as expected due to random fragmentation.

**Figure 4.**
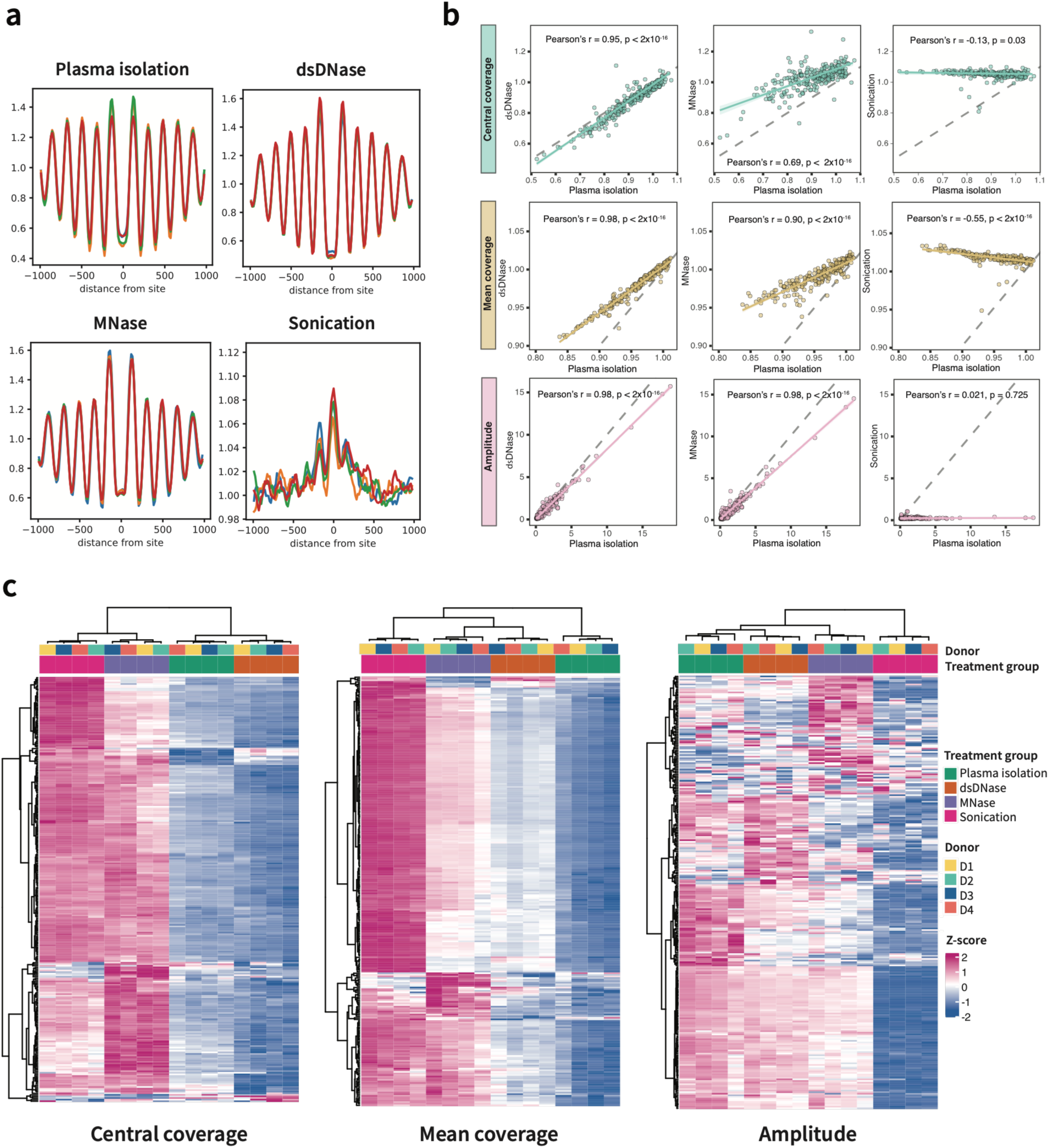
Nucleosome profiles across sample preparation methods. (a) Coverage profiles centered on CTCF binding sites reveal phased nucleosome patterns except for the sonicated samples. (b) Pairwise correlations of nucleosome-derived features, including central coverage, mean coverage, and amplitude, between plasma cfDNA and the three artificial cfDNA samples. Pearson’s and Spearman’s correlation coefficients with corresponding p values are shown in each plot. Each dot represents a TF. Solid lines indicate fitted linear regression trends with confidence intervals, and dashed grey lines indicate the identity line. (c) Heatmaps showing hierarchical clustering of nucleosome-related features across all samples. The columns represent samples annotated by donor and sample treatment group, and the row represent 270 TFs. The values are shown as row-wise z-scores.

We then extracted three nucleosome-derived features, central coverage, mean coverage, and amplitude, to compare each method with plasma-isolated cfDNA. dsDNase and MNase showed strong positive correlations with plasma isolation for most features, indicating partial preservation of nucleosome-associated signals (Figure 4b). Notably, dsDNase-treated DNA revealed the strongest concordance for central coverage (Pearson’s r = 0.95, BH-adjusted p < 0.001), showing only subtle shifts in mean coverage (Pearson’s r was 0.98, BH-adjusted p < 0.001) and amplitude (Pearson’s r = 0.98, BH-adjusted p < 0.001) across all TFs. Sonication showed weak or absent correlation, especially for central coverage and amplitude (Pearson’s r =-0.13 and 0.021, BH-adjusted p = 0.05 and 0.73, respectively), supporting that random fragmentation disrupts nucleosome positioning signals. Hierarchical clustering of nucleosome-derived features further demonstrated that samples were primarily separated by treatment method rather than donor group and that dsDNAse-treated DNA clustered most closely to native cfDNA for 2 of the 3 features (Figure 4c).

### Targeted panel sequencing coverage is affected by fragmentation method

As panel sequencing is currently one of the methods used in clinical practice, we evaluated its performance across the four sample treatment groups. Coverage breadth across the 112 protein-coding genes of the panel was assessed as the number of genes reaching defined depth thresholds, where per-gene mean depth is the average read depth across all target bases of that gene. While clinical panel sequencing typically aims for 10M reads per sample, we obtained an average of 6.25M reads per sample, resulting in lower overall coverage that was nevertheless sufficient to assess differences between the sample types. Up to ≥100x, all 112 genes were covered in every sample, irrespective of sample treatment (Figure 5a, Supplementary Figure 2b). Above ≥200x, differences between groups became apparent: sonicated and dsDNase-treated DNA retained more genes than plasma cfDNA, whereas MNase-treated DNA retained fewer and showed the steepest decline with increasing depth threshold. Analysis of per-gene mean depth similarly showed that MNase-treated DNA generally had the lowest depth, whereas sonicated and dsDNase-treated DNA generally had higher depths (Figure 5b; Supplementary Figures 2a, c, d and 3). Across genes, per-gene mean depth correlated moderately with plasma cfDNA for sonicated and dsDNase-treated DNA (Pearson’s r = 0.75 and 0.66, respectively) and only weakly for MNase-treated DNA (Pearson’s r = 0.33, Figure 5c).

**Figure 5.**
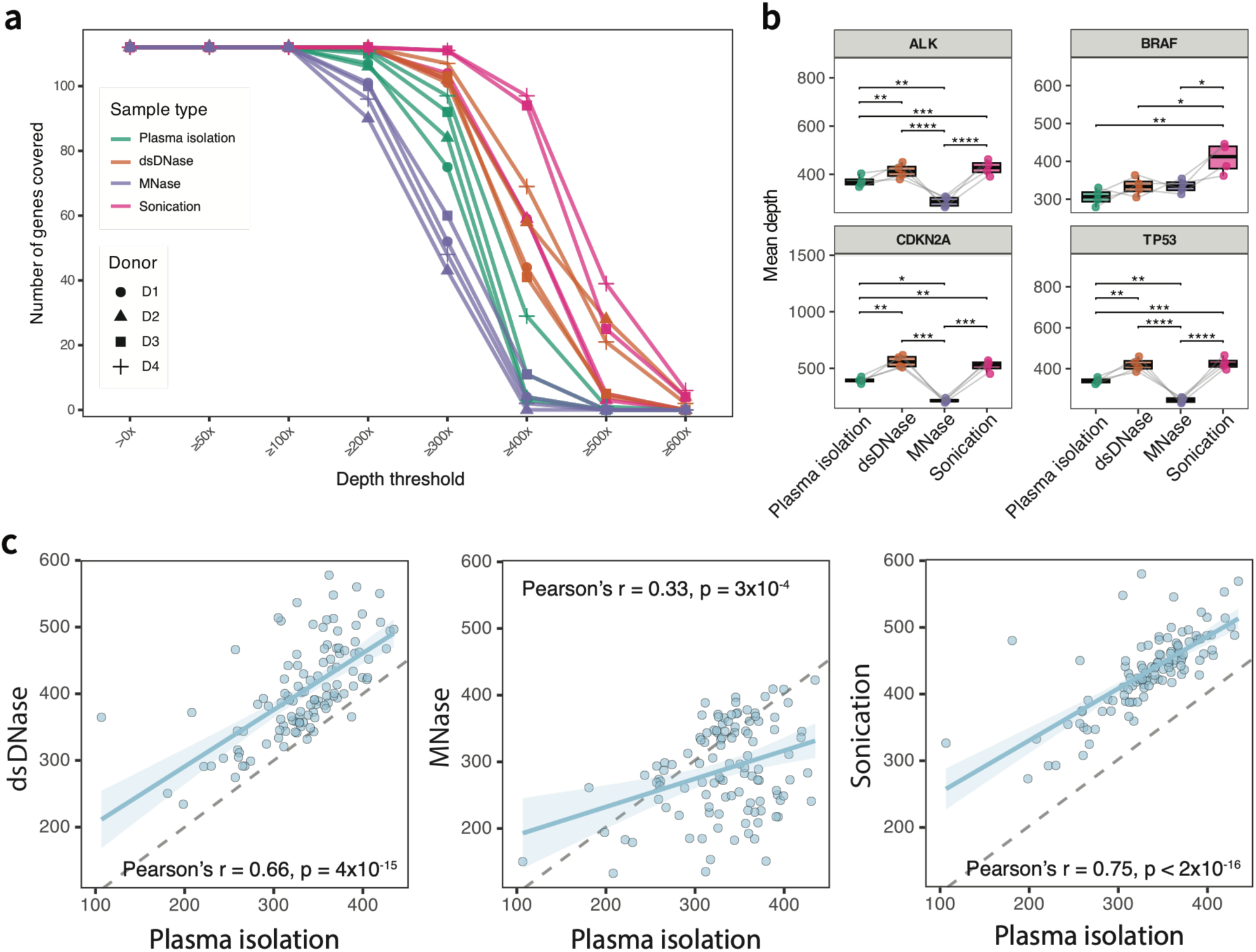
Targeted panel sequencing coverage across 112 protein-coding genes across 3 artificial cfDNA samples and plasma cfDNA. (a) Number of genes covered across increasing depth thresholds for each sample treatment method. Each line represents an individual donor. (b) Mean sequencing depth of four selected genes (*ALK, BRAF, CDKN2A, TP53*). Each point represents one donor, and connecting lines indicate paired donor measurements across sample treatment methods. Statistical significance from Friedman test is indicated as follows: *BH-adjusted p < 0.05, **BH-adjusted p < 0.01, ***BH-adjusted p < 0.001, and ****BH-adjusted p < 0.0001. (c) Pairwise correlations of gene-level mean sequencing depth between plasma isolation and three fragmentation methods. Each dot represents one gene. Pearson’s correlation coefficients with corresponding p values are shown in each plot. Solid lines indicate fitted linear regression trends with confidence intervals, and dashed grey lines indicate the identity line (y=x).

### Tumor/healthy mixtures of dsDNase-based cfDNA better recapitulates tumor specific nucleosome profile compared to MNAse-based cfDNA

To test whether dsDNase and MNase fragmentation retained composition-dependent nucleosome profiles, we mixed artificial cfDNA (dsDNase and MNase) from PBMC with artificial cfDNA from the neuroblastoma CLB-GA cell line and performed sWGS (100% PBMC, 50% PBMC + 50% CLB-GA, and 100% CLB-GA). Tumor fractions estimated using ichorCNA broadly reflected the expected sample compositions for both treatments (Figure 6a and Supplementary Figure 4). Pairwise Manhattan distances from Griffin-derived mean coverage, central coverage, and amplitude profiles across 270 TFs showed consistently lower within-group distances for dsDNase-treated versus MNase-treated PBMC replicates, indicating greater reproducibility of the PBMC nucleosome profiles. In addition, the distances between the two pure sample types (100% PBMC and 100% CLB-GA) were larger after dsDNase treatment for all three features, indicating better discrimination between PBMC-and CLB-GA-derived nucleosome profiles than after MNase treatment (Figure 6b). At the individual TF level, dsDNase treatment better preserved consistent mixture-dependent nucleosome profile features for neuroblastoma-associated TFs, including PHOX2B, HAND2, ASCL1, and GATA3, and PBMC-associated TFs, including SPI1, PAX5, TBX21, and RUNX3, compared with MNase treatment (Figure 6c, d). A similar pattern was also observed for other unselected TFs (Supplementary Figure 5).

**Figure 6.**
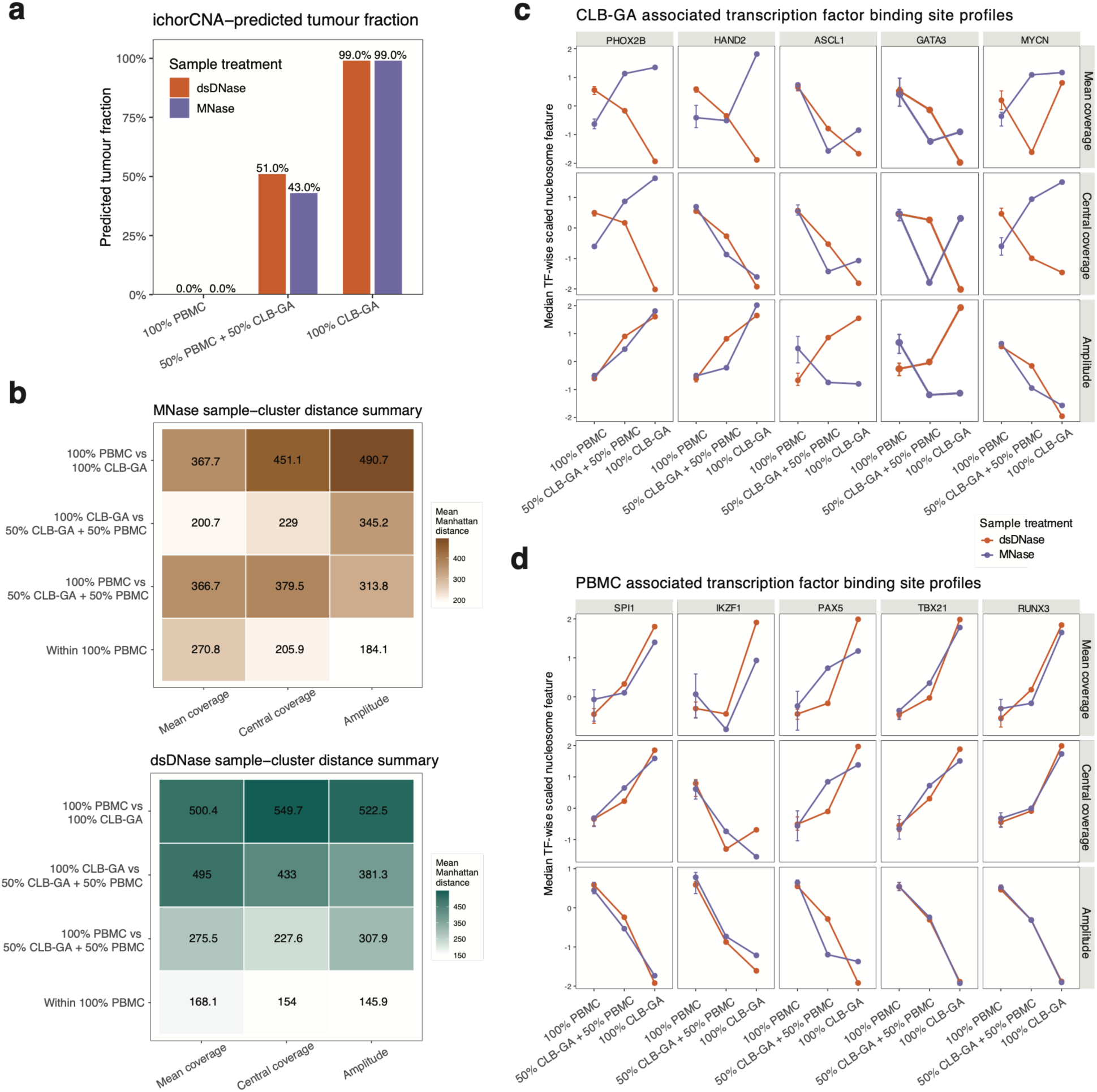
Nucleosome profiles of artificial cfDNA generated by dsDNase and MNase across healthy/tumor mixtures. (a) Tumor fractions in 100% PBMC, 50% PBMC + 50% CLB-GA, and 100% CLB-GA samples were estimated using ichorCNA. (b) Mean pairwise Manhattan distances between tumor/healthy mixtures, calculated from Griffin-derived mean coverage, central coverage, and amplitude, are shown for MNase-treated DNA (top) and dsDNase-treated DNA (bottom). Lower values indicate greater similarity in nucleosome profiles. (c) Mean coverage, central coverage, and amplitude profiles for the neuroblastoma-associated transcription factors PHOX2B, HAND2, ASCL1, GATA3, and MYCN. (d) Corresponding profiles for the PBMC-associated transcription factors SPI1, IKZF1, PAX5, TBX21, and RUNX3.

## Discussion

Benchmarking, validation, and cross-laboratory comparison of cfDNA assay performance are essential for clinical implementation, particularly given the challenge of achieving sufficient sensitivity at low tumor fractions. Representative artificial samples with known aberrations and defined tumor fractions provide a useful way to assess and validate cfDNA assay performance. In this study, we reviewed commercial artificial cfDNA options currently available or under development, each with distinct advantages and limitations. Commercial samples benefit from optimized and standardized workflows, ensuring reproducibility, but are generally more costly than laboratory-generated reference materials. Most systems involve synthetic DNA spiked into a healthy background, offering flexibility to introduce specific genomic aberrations, but limiting utility to a restricted set of variants or cancer types. These references are also not suitable for epigenomic assays, including cfDNA methylation profiling or fragmentomics. Recently, some companies have begun offering artificial cfDNA with mononucleosomal and dinucleosomal fragments, though these remain limited to specific cell lines or common cancer types. For rarer tumor types, more flexible and biologically representative reference material is still needed.

We compared three workflows for generating artificial cfDNA: two enzymatic fragmentation methods (MNase and dsDNase) and one mechanical method (sonication) to determine which approach yields the most representative plasma cfDNA sample. As expected, sonicated samples lacked several characteristic features of native plasma cfDNA, including fragment size pattern and nucleosome footprints due to random fragmentation. In contrast, enzymatic fragmentation more closely mimicked apoptotic processes that generate cfDNA *in vivo*. Fragment length profiles using Tapestation confirm that enzymatic digestion produced fragments with characteristic length distribution for cfDNA. However, enzymatically fragmented DNA showed an approximately 10 bp shorter fragment length distribution compared with plasma cfDNA, with higher abundance of short fragments in the 50–150 bp range and more pronounced 10 bp periodicity. Following library preparation, dsDNase-treated samples showed an additional shift toward shorter insert sizes. This is likely explained by the prominent single-stranded overhangs generated by dsDNase, which are subject to exonucleolytic trimming during library preparation, resulting in shorter final library inserts^60^. A similar shift toward shorter insert sizes was also observed in sonicated samples. One possible explanation is that sonication generates heterogeneous and chemically abnormal DNA termini that may be inefficiently processed during end repair and adapter ligation^61^. Sonication may also introduce internal strand damage; if such damage is not repaired and disrupts the continuity of both strands within an adapter-flanked molecule, recovery of the full length insert during library amplification may be reduced. Preferential loss of these molecules could therefore contribute to the shorter final insert size distribution, although this mechanism was not directly investigated in our study^62^. Nucleosome footprinting via Griffin demonstrated that enzymatically digested DNA retained nucleosome occupancy patterns typical for plasma cfDNA. Particularly, dsDNase digestion showed greater similarity to plasma cfDNA nucleosomal patterns than MNase fragmentation. These findings support the use of enzymatic fragmentation when preservation of nucleosome-associated fragmentomic information is important.

Targeted panel sequencing showed that fragmentation method affected gene-level sequencing coverage. Across most genes, MNase-treated DNA showed the lowest per-gene mean depth, whereas dsDNase-treated and sonicated DNA generally showed higher depths. These coverage differences are unlikely to substantially affect the detection and variant allele fraction (VAF) estimation of variants present at relatively high VAFs, for which sufficient variant-supporting reads are expected. However, at low tumor fractions and low VAFs, reduced coverage may decrease the number of variant-supporting reads, thereby reducing detection sensitivity and increasing uncertainty in VAF estimates. Analytical comparisons between artificial DNA and plasma cfDNA should therefore account for locus-specific coverage and preferably focus on variant loci with sufficient coverage across sample types.

Based on fragment size distribution, nucleosome occupancy and panel-sequencing coverage analyses, enzymatically fragmented DNA, particularly dsDNase-treated DNA, most closely resembles native cfDNA. However, for detailed fragmentomic applications, including end-motif analysis neither enzymatic nor sonication approaches fully recapitulated native cfDNA features. This limitation reflects how DNA end motifs are not merely technical features but reflect nuclease activity and biological cfDNA generation processes that are inherently tied to the fragmentation mechanism, including the specific nuclease or physical shearing process. Enzymatic fragmentation using dsDNase I or MNase will therefore generate end-motif signatures that differ from native plasma cfDNA, which is shaped *in vivo* by endogenous nucleases, including DNase1L3^63,64^. Consequently, artificial fragmentation can introduce method-specific cleavage signatures that may confound fragmentomics-based biomarker analyses.

There is also a trade-off between biological representation and scalability. Generating enzymatically fragmented DNA requires live cell input and approximately four hours of hands-on work. Enzyme concentrations often need to be optimized for each cell line, and small laboratory handing variations can lead to over-or under-digestion, frequently requiring repeat experiments. Rigorous quality control is essential, with fragment length profiles serving as a key metric: the first nucleosome peak should be around 160-170 bp, and the sample should show nucleosomal peaks without a high molecular weight peak. In contrast, sonication is well suited for high-throughput experiments, enabling the production of large amounts of DNA in a straightforward workflow requiring minimal optimization. For genomic assays detecting SNVs, indels and CNVs, sonicated artificial cfDNA will be a suitable model for benchmarking studies. However, as the field moves towards clinical implementation of fragmentomics-based assays, the need for artificial cfDNA models preserving biological fragmentation information will increase.

In conclusion, enzymatic fragmentation preserved nucleosome-associated organization and cfDNA-like fragment-length profiles more effectively than sonication, with dsDNase treatment providing the closest overall resemblance to plasma cfDNA. Still, enzyme fragmented DNA remains an imperfect model, with enzyme specific cleavage biases limiting detailed fragmentomic applications, and additional hands-on time, protocol optimization, and sensitivity to digestion conditions constraining scalability.

## Materials and methods

### Blood collection and plasma isolation

The study was approved by the ethical committee and informed consent was obtained. Donors declared themselves healthy at blood collection. Six 10 ml K2E EDTA tubes were collected per donor. Plasma was isolated by double centrifugation: 1900 g for 10 minutes, transfer of platelet poor plasma without disturbing the buffy coat, and a second 1900 g for 10 minutes spin to obtain platelet free plasma. Plasma was stored at −80°C until cfDNA extraction. cfDNA was extracted using the Maxwell RSC LV ccfDNA kit (Promega) according to protocol for large volumes.

### Cell culture

ALK mutated neuroblastoma CLB-GA cells were cultured in RPMI-1640 supplemented with 10 % FCS, 2 mM L-glutamine, and 25 mM HEPES at 37°C and 5 % CO2. Mycoplasma testing (Lonza) and STR genotyping were performed regularly. For enzymatic fragmentation, cells were used immediately after harvesting or frozen in 10 % DMSO and stored at −80°C.

### PBMC isolation

Peripheral blood mononuclear cells (PBMCs) were isolated using Leucosep tubes (50 ml; 227289, Greiner) prepared with 15 ml Ficoll Paque Plus (GE healthcare) and centrifuged 30 seconds at 1000 g. Buffy coat was diluted with PBS (1X, Gibco) and added to the tube, followed by centrifugation of 18 minutes at 800 g, brakes off. The enriched fraction was washed with 5 ml PBS and centrifuged 10 minutes at 250 g at 3°C. Red blood cells were lysed with ACK buffer (Gibco) for 5 minutes, stopped by PBS dilution, and washed again. Cells were counted and aliquoted in 50 percent PBS/FCS with 10 percent DMSO, frozen in a freezing container, and stored at −80°C.

### Preparation of enzymatically fragmented and sonicated samples

Enzymatic fragmentation was performed according to the supplier protocol of EZ Nucleosomal DNA prep kit (Zymo). PBMCs and CLB-GA cells were digested with 0.1 U or 1 U MNase, and 0.7 U or 1.2 U dsDNase, for PBMC samples and tumor cell culture samples respectively.

Genomic DNA from PBMCs was isolated using the Qiamp DNA mini and Blood mini kit (Qiagen) and eluted in nuclease free water. One microgram DNA in 100 µl 0.1x TE was sonicated on ice for 15 minutes using a Bioruptor at low intensity for 70 cycles (30 ON, 30 OFF). Fragment length was assessed using the cell-free DNA screentape on Agilent TapeStation and compared to the length profile of plasma derived cfDNA; samples that were more than 30 bp longer than native cfDNA received additional 20 cycle sonication blocks.

### Shallow whole-genome sequencing and preprocessing

Libraries were prepared using NEXTflex Cell Free DNA Seq kit. Sixteen libraries were pooled and sequenced on an Illumina NextSeq 2000 (P3 XLEAP SBS, 100 cycles) targeting 75 million reads per sample. FASTQ conversion and demultiplexing used nf-core/demultiplex^48^. Trim Galore removed adapters and low-quality bases. Reads were aligned to GRCh38 with Bowtie2, sorted with SAMtools, and deduplicated with UMI tools.

### Fragmentomic extraction

Fragmentomic features were extracted following Liu et al.^49^ BAM files were filtered for properly paired reads, excluding unmapped reads, duplicates, secondary alignments, and reads with mapping quality below 30. GC content was calculated for GC bias correction. Nucleosome profiling used Griffin^50^ with 30000 transcription factor binding sites (TFBSs) for 270 transcription factors (TFs) from GTRD^51^. Fragments with lengths between 100 and 200 bp were retained. GC corrected midpoint coverage profiles were generated, and three nucleosome features were extracted: central coverage, mean coverage, and peak amplitude.

### Panel sequencing coverage analysis

Panel sequencing using KAPA HyperCap (Roche) with a custom 112 gene tumor panel was performed on libraries generated from sWGS as described above. The table of genes included within the panel is listed in supplementary table 1. Enriched libraries were sequenced on a NextSeq 2000 (P1 XLEAP SBS, 100 cycles) yielding an average of 6.25 million reads per sample. Data was processed similar as sWGS. Coverage analysis for 112 genes used mosdepth^52^. BAM files were downsampled to match the lowest read count. For each gene, mean depth and proportions of bases above 100x, 200x, 300x, 500x, and 600x were extracted.

## Statistical analysis

Analyses were performed in R (version 2025.09.2+418). Donor matched groups (plasma, dsDNase, MNase, sonication) were compared using the Friedman test with Benjamini–Hochberg (BH) false discovery rate correction for multiple testing. Correlations used Pearson’s correlation coefficient for approximately normal variables and Spearman’s rank correlation coefficient (Spearman’s ρ) for non-parametric variables.

## Acknowledgements

We gratefully acknowledge the non-invasive prenatal testing (NIPT) team at the Center for Medical Genetics Ghent (CMGG) for performing the shallow whole-genome library preparation.

## Author contributions

K.D.P. conceptualized and supervised the study and wrote the manuscript.

L.C. conceptualized the study, performed laboratory experiments and data analysis, interpretation and visualization, and wrote the manuscript.

T.N.N. performed data analysis, interpretation and visualization, and wrote the manuscript.

R.V.B., S.R., A.D.C. and J.V.D.M. performed laboratory experiments

K.D.P., L.C., T.N.N., A.D.C., J.V.D.M., and N.V.R. reviewed and edited the manuscript.

