## Supplementary materials for "Benchmarking fragmentation-derived artificial cfDNA reference standards"

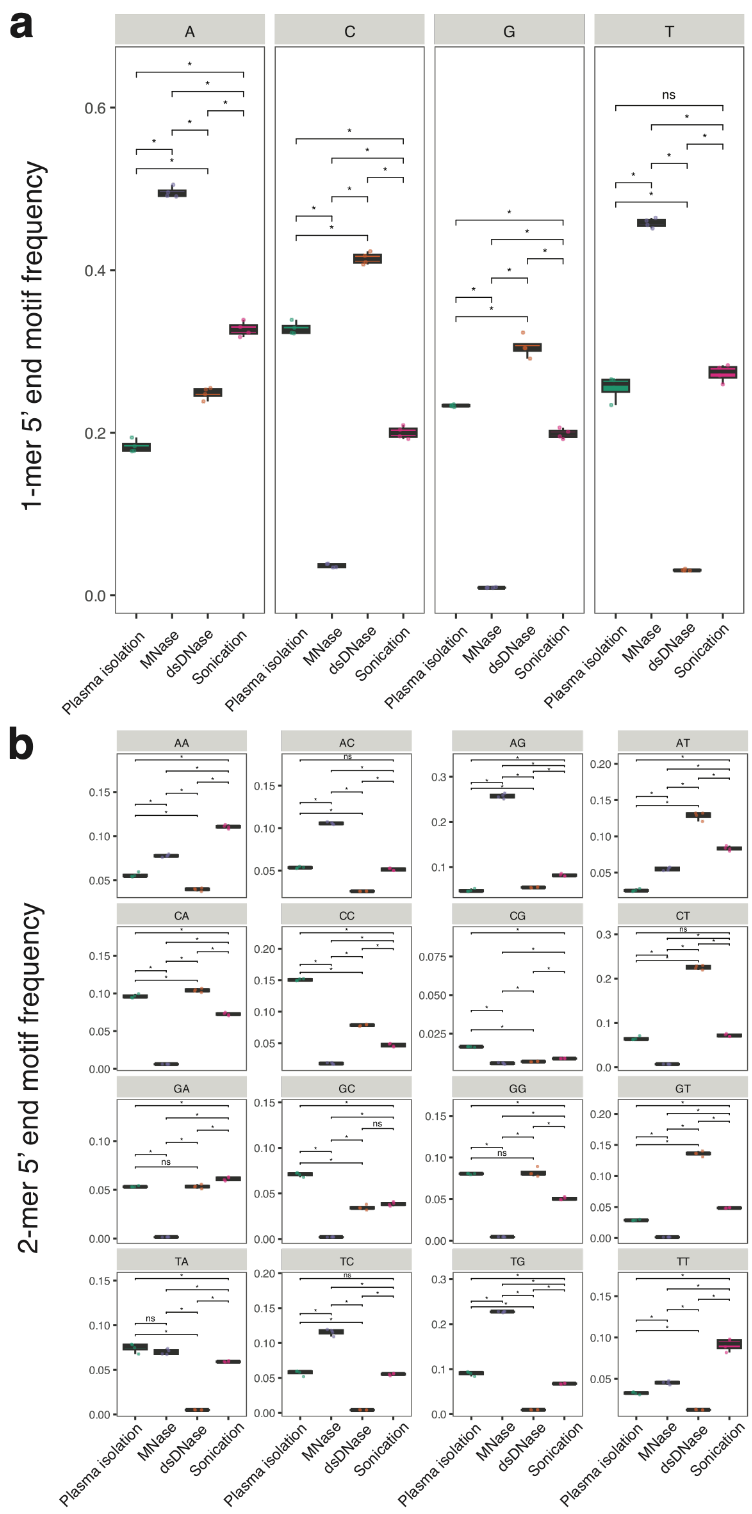


***Supplementary Figure 1.*** 5' end-motif frequencies differ across plasma cfDNA, MNase digested, dsDNase digested, and sonicated DNA. (a) Frequencies of 1-mer 5' end motifs. (b) Frequencies of 2-mer 5' end motifs. (*) Friedman test, BH-adjusted $p < 0.05$; ns: not significant.

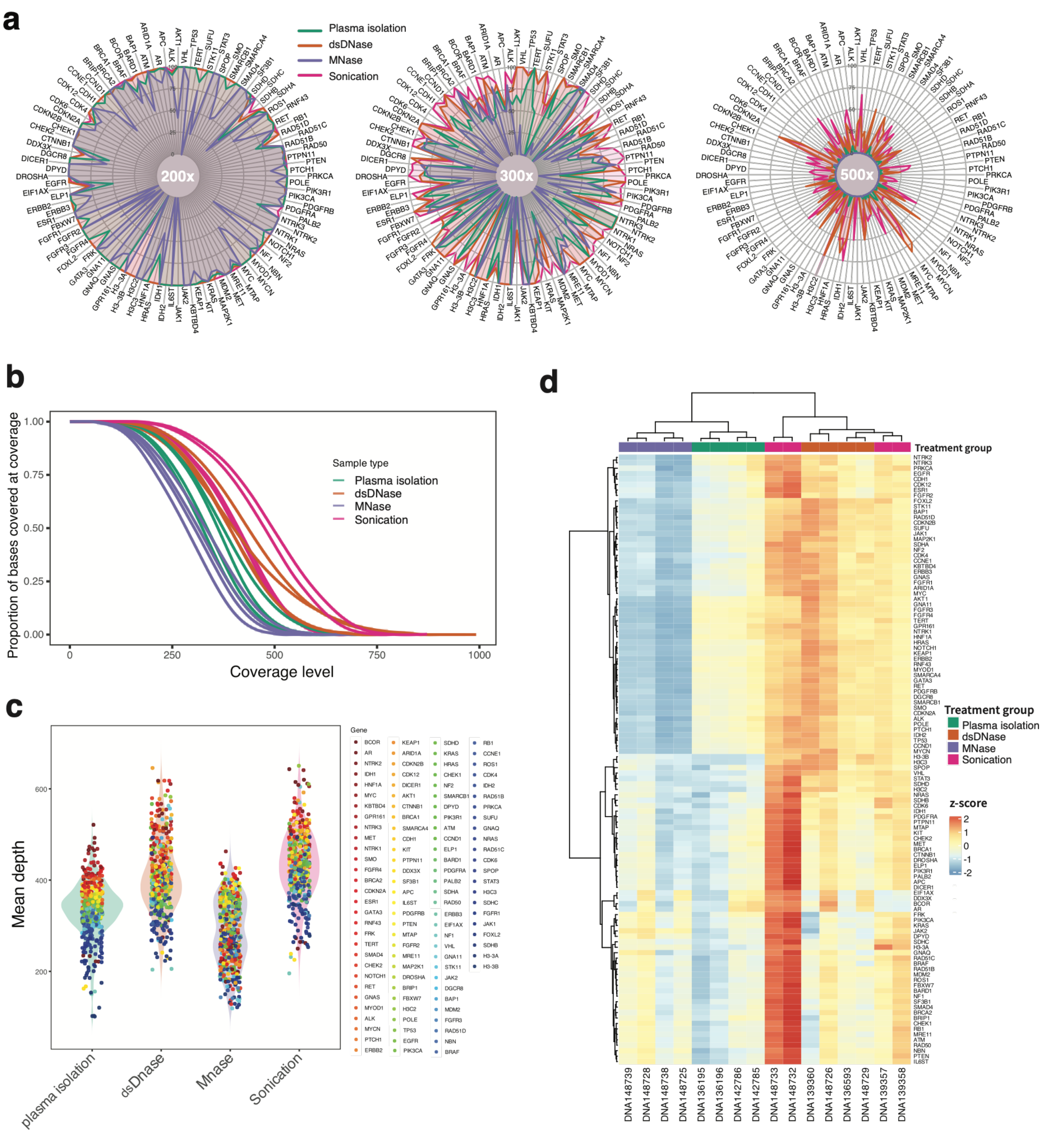


***Supplementary Figure 2.*** (a) Radar plots show gene-level coverage across all 112 genes at three depth thresholds: 200x, 300x, and 500x. Each axis corresponds to one gene in the panel, and lines represent four sample treatment methods. (b) The proportion of target bases covered at increasing sequencing depth thresholds across sample treatment groups, including plasma cfDNA, dsDNase-treated DNA, MNase-treated DNA, and sonicated DNA. (c) Distribution of gene-level mean depth across treatment groups. Each point represents one targeted gene, and colors denote individual genes. Violin plots summarize the overall coverage distribution for each sample treatment method. (d) Unsupervised hierarchical clustering heatmap of gene-level mean coverage across samples. Coverage values were standardized by gene and displayed as z-scores.

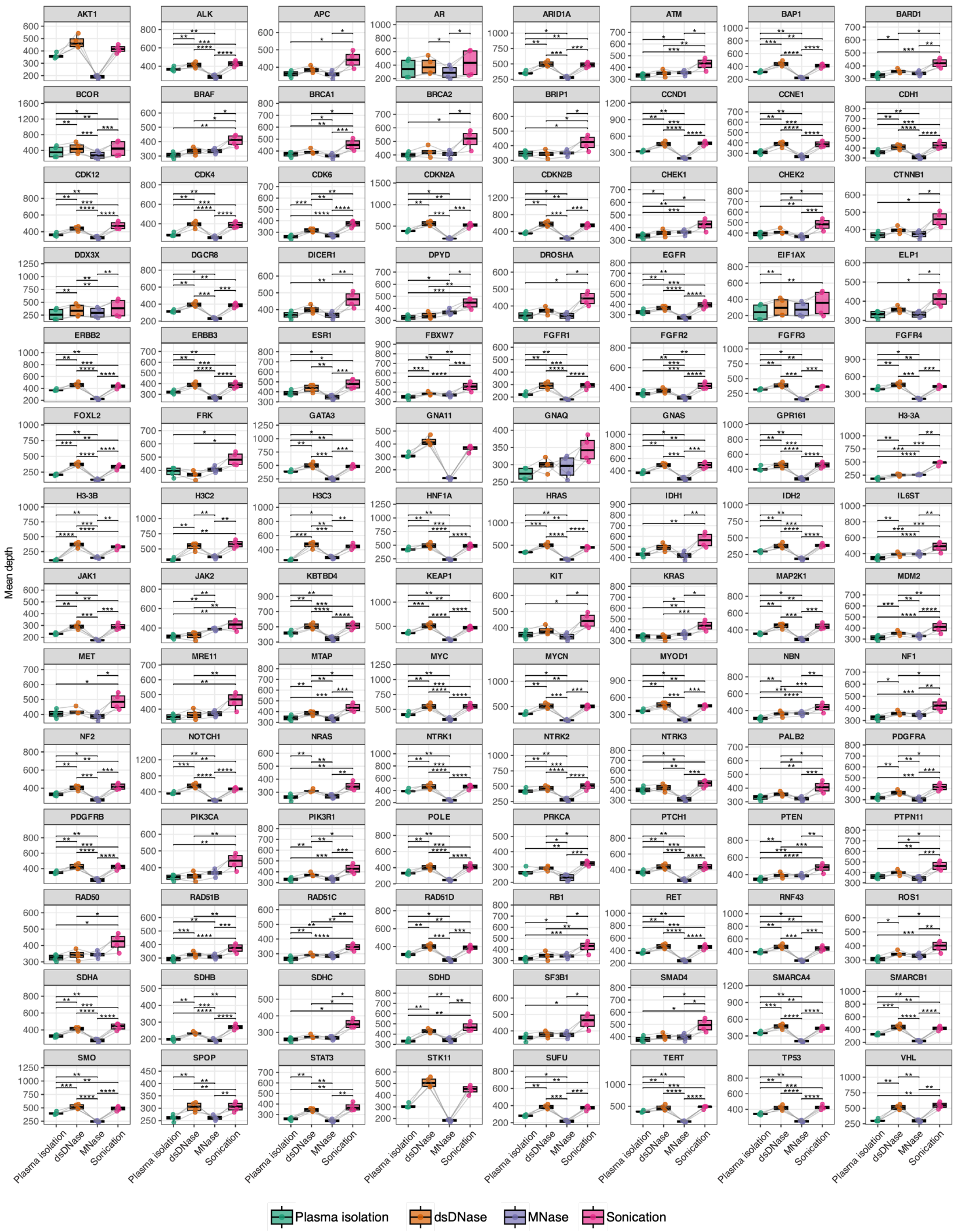


***Supplementary Figure 3.*** Gene-specific coverage differences across sample treatment methods. Each point represents one donor, and connecting lines indicate paired donor measurements across sample treatment methods. Statistical significance from Friedman test is indicated as follows: *BH-adjusted $p < 0.05$, **BH-adjusted $p < 0.01$, ***BH-adjusted $p < 0.001$, and ****BH-adjusted $p < 0.0001$.

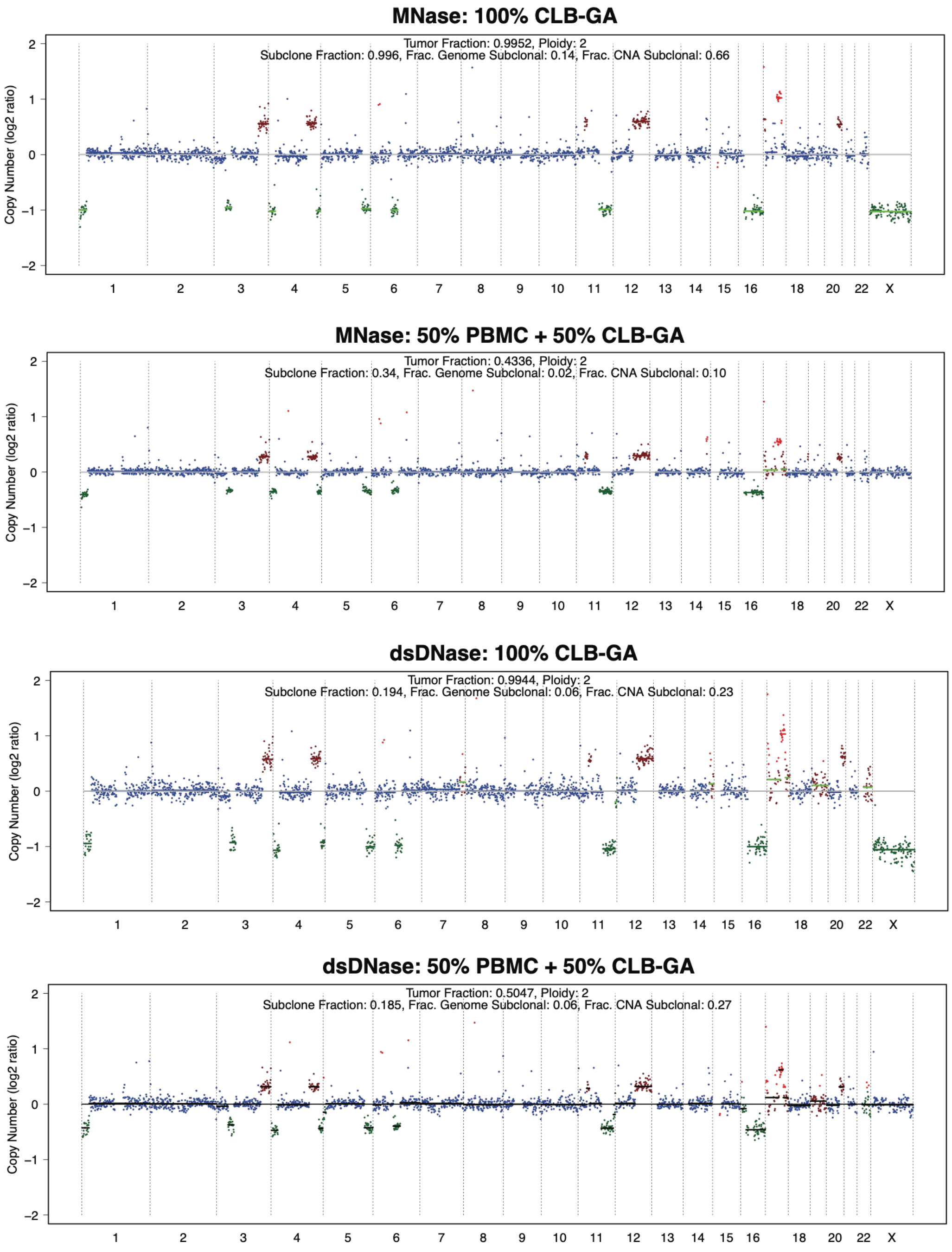


***Supplementary Figure 4.*** Genome wide copy number profiles generated using ichorCNA for 100% CLB-GA and 50% PBMC + 50% CLB-GA samples following MNase treatment (top two panels) or dsDNase treatment (bottom two panels). Copy number log2 ratios are shown across the autosomes and chromosome X. The corresponding ichorCNA estimates of tumor fraction were 99.5% and 43.4% for the MNase-treated 100% CLB-GA and 50:50 mixture samples, respectively, and 99.4% and 50.5% for the corresponding dsDNase-treated samples.

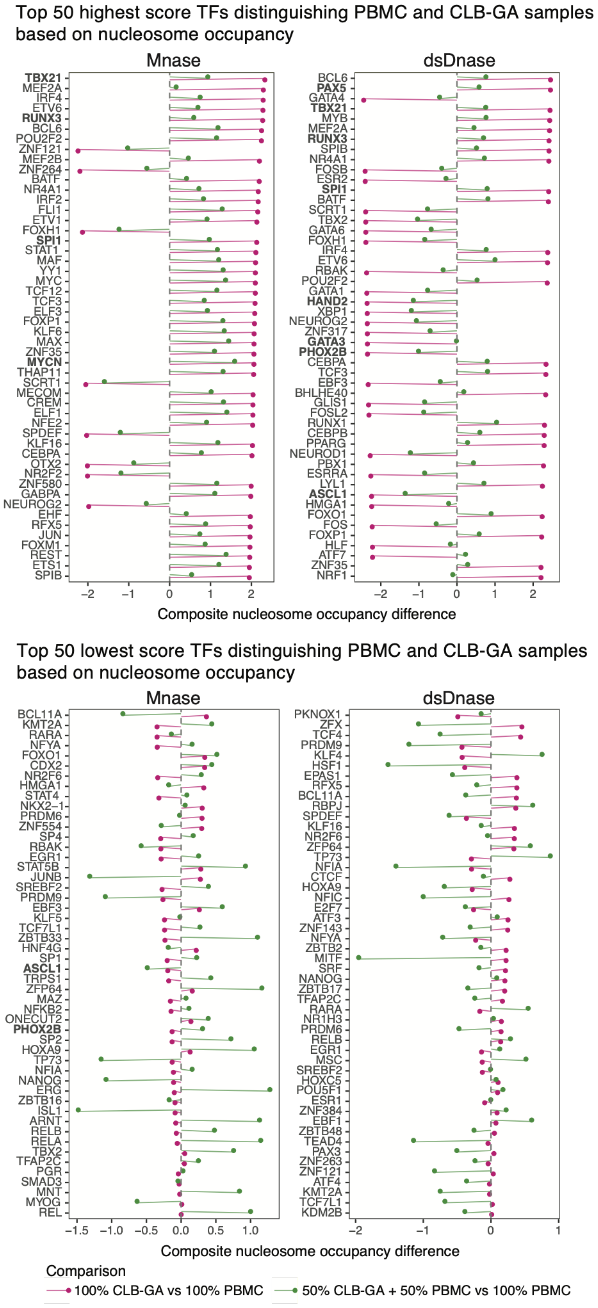


***Supplementary Figure 5.*** **Transcription factors with the largest differences in composite nucleosome occupancy between PBMC- and CLB-GA-derived artificial cfDNA after MNase or dsDNase fragmentation.** (**a**) The 50 transcription factors (TFs) with the highest and (**b**) the 50 with the lowest composite nucleosome occupancy difference, ranked separately for each fragmentation method. For each TF, the composite score integrates Griffin-derived mean coverage, central coverage, and amplitude across its associated binding sites; plotted values are the difference in this score between a CLB-GA-containing sample and the 100% PBMC sample. Two comparisons are shown: 100% CLB-GA versus 100% PBMC, and 50% CLB-GA + 50% PBMC versus 100% PBMC. Positive values indicate greater composite nucleosome occupancy in the CLB-GA-containing sample, negative values lower occupancy.

### Supplementary Table 1. Composition of the targeted sequencing panel.

The panel comprises 112 genes (70 sequenced across the full coding region, 42 covered at selected hotspots only), corresponding to 1172 target regions. For each gene, the reference transcript is given as the NCBI RefSeq and Ensembl (release 105) identifier. Target regions are listed as chromosomal coordinates (chr:start-end) on the GRCh38/hg38 assembly and are separated by commas.

| No. | Gene | Panel coverage | RefSeq transcript (NCBI) | Ensembl transcript ID | Target regions (GRCh38/hg38) |
|---|---|---|---|---|---|
| 1 | ***AKT1*** | Hotspot | NM_005163.2 | ENST00000555528 | chr14:104780088-104780216, chr14:104776659-104776770, chr14:104772878-104773092 |
| 2 | ***ALK*** | Hotspot | NM_004304.5 | ENST00000389048 | chr2:29225461-29225565, chr2:29223342-29223528, chr2:29222517-29222607, chr2:29222344-29222408, chr2:29220706-29220835, chr2:29213984-29214081, chr2:29209786-29209878, chr2:29207171-29207272, chr2:29197542-29197676, chr2:29196770-29196860, chr2:29193223-29193922 |
| 3 | ***APC*** | Full gene | NM_000038.6 | ENST00000257430 | chr5:112754891-112755025, chr5:112766326-112766410, chr5:112767189-112767390, chr5:112775629-112775737, chr5:112780790-112780903, chr5:112792446-112792529, chr5:112801279-112801383, chr5:112815495-112815593, chr5:112818966-112819344, chr5:112821896-112821991, chr5:112827108-112827247, chr5:112827929-112828006, chr5:112828856-112828972, chr5:112834951-112835165, chr5:112837553-112844127 |
| 4 | ***AR*** | Hotspot | NM_000044.6 | ENST00000374690 | chrX:67711402-67711689, chrX:67717478-67717622, chrX:67723686-67723841 |
| 5 | ***ARID1A*** | Full gene | NM_006015.6 | ENST00000324856 | chr1:26696402-26697540, chr1:26729651-26729863, chr1:26731152-26731604, chr1:26732676-26732792, chr1:26760856-26761096, chr1:26761384-26761473, chr1:26762152-26762319, chr1:26762973-26763285, chr1:26766221-26766366, chr1:26766457-26766566, chr1:26767790-26767999, chr1:26771119-26771326, chr1:26772500-26772632, chr1:26772812-26772987, chr1:26773346-26773496, chr1:26773580-26773717, chr1:26773802-26773898, chr1:26774329-26775220, chr1:26775577-26775707, chr1:26779023-26780758 |
| 6 | ***ATM*** | Full gene | NM_000051.4 | ENST00000675843 | chr11:108227624-108227696, chr11:108227775-108227888, chr11:108229177-108229323, chr11:108235669-108235834, chr11:108243952-108244118, chr11:108244787-108245026, chr11:108246963-108247127, chr11:108248932-108249102, chr11:108250700-108251072, chr11:108251836-108252031, chr11:108252816-108252912, chr11:108253813-108254039, chr11:108256214-108256340, chr11:108257480-108257606, chr11:108258985-108259075, chr11:108267170-108267342, chr11:108268409-108268609, chr11:108271063-108271146, chr11:108271250-108271406, chr11:108272531-108272607, chr11:108272721-108272852, chr11:108279490-108279608, chr11:108280994-108281168, chr11:108282709-108282879, chr11:108284226-108284473, chr11:108287599-108287715, chr11:108288976-108289103, chr11:108289601-108289801, chr11:108292618-108292793, chr11:108293312-108293477, chr11:108294926-108295059, chr11:108297286-108297382, chr11:108299713-108299885, chr11:108301647-108301789, chr11:108302852-108303029, chr11:108304674-108304852, chr11:108307896-108307984, chr11:108310159-108310315, chr11:108312410-108312498, chr11:108315822-108315911, chr11:108316010-108316113, chr11:108317372-108317521, chr11:108319953-108320058, chr11:108321300-108321420, chr11:108325309-108325544, chr11:108326057-108326225, chr11:108327644-108327758, chr11:108329020-108329238, chr11:108330213-108330421, chr11:108331443-108331557, chr11:108331878-108332037, chr11:108332761-108332900, chr11:108333885-108333968, chr11:108334968-108335109, chr11:108335844-108335961, chr11:108343221-108343371, chr11:108345742-108345908, chr11:108347278-108347365, chr11:108353765-108353880, chr11:108354810-108354874, chr11:108365081-108365218, chr11:108365324-108365508 |
| 7 | ***BAP1*** | Full gene | NM_004656.4 | ENST00000460680 | chr3:52409842-52409880, chr3:52409714-52409743, chr3:52409554-52409608, chr3:52408474-52408606, chr3:52407958-52408077, chr3:52407399-52407460, chr3:52407174-52407316, chr3:52406829-52406907, chr3:52406253-52406376, chr3:52405765-52405912, chr3:52405110-52405294, chr3:52404453-52404586, chr3:52403416-52403894, chr3:52403138-52403298, chr3:52402779-52402871, chr3:52402602-52402674, chr3:52402286-52402421 |
| 8 | ***BARD1*** | Full gene | NM_000465.4 | ENST00000260947 | chr2:214809412-214809571, chr2:214797061-214797117, chr2:214792297-214792445, chr2:214780560-214781509, chr2:214769232-214769312, chr2:214767482-214767654, chr2:214752447-214752555, chr2:214745722-214745854, chr2:214745067-214745159, chr2:214730411-214730508, chr2:214728674-214729008 |
| 9 | ***BCOR*** | Full gene | NM_001123385.2 | ENST00000378444 | chrX:40077844-40077930, chrX:40076454-40076532, chrX:40072349-40075180, chrX:40071637-40071690, chrX:40070973-40071159, chrX:40064336-40064599, chrX:40063608-40063952, chrX:40062746-40063071, chrX:40062139-40062393, chrX:40057155-40057321, chrX:40055368-40055513, chrX:40054256-40054333, chrX:40053886-40054042, chrX:40052107-40052400 |
| 10 | ***BRAF*** | Hotspot | NM_004333.6 | ENST00000646891 | chr7:140801412-140801560, chr7:140800362-140800481, chr7:140781576-140781693, chr7:140777991-140778075, chr7:140754187-140754233, chr7:140753275-140753393 |
| 11 | ***BRCA1*** | Full gene | NM_007294.4 | ENST00000357654 | chr17:43125251-43125384, chr17:43123997-43124135, chr17:43115706-43115799, chr17:43106436-43106553, chr17:43104848-43104976, chr17:43104102-43104281, chr17:43099755-43099900, chr17:43097224-43097309, chr17:43095826-43095942, chr17:43091415-43094880, chr17:43090924-43091052, chr17:43082384-43082595, chr17:43076468-43076634, chr17:43074311-43074541, chr17:43070908-43071258, chr17:43067588-43067715, chr17:43063854-43063971, chr17:43063313-43063393, chr17:43057032-43057155, chr17:43051043-43051137, chr17:43049101-43049214, chr17:43047623-43047723, chr17:43045656-43045822 |
| 12 | ***BRCA2*** | Full gene | NM_000059.4 | ENST00000380152 | chr13:32315454-32315687, chr13:32316402-32316547, chr13:32319057-32319345, chr13:32325056-32325204, chr13:32326081-32326170, chr13:32326222-32326302, chr13:32326479-32326633, chr13:32329423-32329512, chr13:32330899-32331050, chr13:32332252-32333407, chr13:32336245-32341216, chr13:32344538-32344673, chr13:32346807-32346916, chr13:32354841-32355308, chr13:32356408-32356629, chr13:32357722-32357949, chr13:32362503-32362713, chr13:32363159-32363553, chr13:32370382-32370577, chr13:32370936-32371120, chr13:32376650-32376811, chr13:32379297-32379535, chr13:32379730-32379933, chr13:32379987-32380165, chr13:32394669-32394953, chr13:32396878-32397064, chr13:32398142-32398792 |
| 13 | ***BRIP1*** | Full gene | NM_032043.3 | ENST00000259008 | chr17:61861447-61861541, chr17:61859796-61859907, chr17:61857058-61857231, chr17:61849129-61849256, chr17:61847101-61847220, chr17:61808467-61808757, chr17:61801253-61801474, chr17:61799100-61799299, chr17:61793597-61793729, chr17:61784270-61784424, chr17:61780840-61781005, chr17:61780261-61780401, chr17:61776401-61776562, chr17:61744432-61744591, chr17:61743013-61743134, chr17:61715951-61716063, chr17:61693430-61693512, chr17:61685836-61686165, chr17:61683294-61684140 |
| 14 | ***CCND1*** | Full gene | NM_053056.3 | ENST00000227507 | chr11:69641314-69641511, chr11:69643031-69643246, chr11:69643832-69643991, chr11:69647994-69648142, chr11:69651118-69651282 |
| 15 | ***CCNE1*** | Full gene | NM_001238.4 | ENST00000262643 | chr19:29812555-29812578, chr19:29812689-29812776, chr19:29812969-29813037, chr19:29817137-29817282, chr19:29817406-29817541, chr19:29820702-29820848, chr19:29821722-29821817, chr19:29821996-29822130, chr19:29822240-29822351, chr19:29822446-29822603, chr19:29823655-29823779 |
| 16 | ***CDH1*** | Full gene | NM_004360.5 | ENST00000261769 | chr16:68737415-68737463, chr16:68738297-68738411, chr16:68801670-68801893, chr16:68808424-68808567, chr16:68808693-68808848, chr16:68810197-68810341, chr16:68811684-68811859, chr16:68812135-68812263, chr16:68813313-68813495, chr16:68815515-68815759, chr16:68819280-68819425, chr16:68822001-68822225, chr16:68823399-68823626, chr16:68828174-68828304, chr16:68829654-68829797, chr16:68833290-68833501 |
| 17 | ***CDK4*** | Full gene | NM_000075.4 | ENST00000257904 | chr12:57751500-57751717, chr12:57751207-57751342, chr12:57750923-57751090, chr12:57750656-57750765, chr12:57749454-57749504, chr12:57749182-57749317, chr12:57748525-57748617 |
| 18 | ***CDK6*** | Full gene | NM_001259.8 | ENST00000265734 | chr7:92833091-92833324, chr7:92774696-92774831, chr7:92725626-92725793, chr7:92671426-92671535, chr7:92623036-92623086, chr7:92618072-92618207, chr7:92615139-92615286 |
| 19 | ***CDK12*** | Full gene | NM_016507.4 | ENST00000447079 | chr17:39462071-39463117, chr17:39470879-39471763, chr17:39490557-39490733, chr17:39492751-39492890, chr17:39494524-39494694, chr17:39501250-39501439, chr17:39509705-39509761, chr17:39511529-39511630, chr17:39515731-39515808, chr17:39517440-39517556, chr17:39519956-39520087, chr17:39524674-39524885, chr17:39525864-39526316, chr17:39530604-39531318 |
| 20 | ***CDKN2A*** | Full gene | NM_000077.5 | ENST00000304494 | chr9:21974678-21974827, chr9:21970902-21971208, chr9:21968229-21968242 |
| 21 | ***CDKN2B*** | Full gene | NM_004936.4 | ENST00000276925 | chr9:22008798-22008953, chr9:22005987-22006247 |
| 22 | ***CHEK1*** | Full gene | NM_001114122.3 | ENST00000438015 | chr11:125626768-125626833, chr11:125627607-125627830, chr11:125629232-125629296, chr11:125629391-125629460, chr11:125633163-125633351, chr11:125635429-125635533, chr11:125637449-125637544, chr11:125643792-125643900, chr11:125644091-125644268, chr11:125644512-125644643, chr11:125653746-125653847, chr11:125655225-125655322 |
| 23 | ***CHEK2*** | Full gene | NM_007194.4 | ENST00000404276 | chr22:28734403-28734721, chr22:28725243-28725367, chr22:28724977-28725124, chr22:28719395-28719485, chr22:28711909-28712017, chr22:28710006-28710059, chr22:28703505-28703566, chr22:28699838-28699937, chr22:28696901-28696987, chr22:28695710-28695873, chr22:28695127-28695242, chr22:28694032-28694117, chr22:28689135-28689215, chr22:28687895-28687986 |
| 24 | ***CTNNB1*** | Hotspot | NM_001904.4 | ENST00000349496 | chr3:41224067-41224081, chr3:41224526-41224753, chr3:41224954-41225207, chr3:41227208-41227352, chr3:41233341-41233444 |

| No. | Gene | Panel coverage | RefSeq transcript (NCBI) | Ensembl transcript ID | Target regions (GRCh38/hg38) |
|---|---|---|---|---|---|
| 25 | ***DGCR8*** | Full gene | NM_022720.7 | ENST00000351989 | chr22:20085963-20086683, chr22:20087162-20087321, chr22:20089669-20089811, chr22:20089976-20090258, chr22:20091435-20091632, chr22:20091869-20091970, chr22:20092809-20092907, chr22:20094713-20094795, chr22:20106177-20106277, chr22:20106592-20106698, chr22:20107271-20107398, chr22:20108890-20109003, chr22:20110025-20110109 |
| 26 | ***DDX3X*** | Full gene | NM_001356.5 | ENST00000644876 | chrX:41334252-41334297, chrX:41337408-41337465, chrX:41339036-41339083, chrX:41341484-41341616, chrX:41342495-41342653, chrX:41342737-41342836, chrX:41343216-41343351, chrX:41343737-41343822, chrX:41344030-41344128, chrX:41344239-41344399, chrX:41345180-41345324, chrX:41345404-41345548, chrX:41346229-41346410, chrX:41346505-41346622, chrX:41346859-41347012, chrX:41347312-41347451, chrX:41347640-41347720 |
| 27 | ***DICER1*** | Full gene | NM_030621.4 | ENST00000526495 | chr14:95133315-95133459, chr14:95132515-95132677, chr14:95131509-95131639, chr14:95130058-95130192, chr14:95129472-95129632, chr14:95126580-95126748, chr14:95124196-95124668, chr14:95117622-95117754, chr14:95116453-95116695, chr14:95115667-95115821, chr14:95113092-95113224, chr14:95112172-95112247, chr14:95111317-95111456, chr14:95108324-95108503, chr14:95107880-95108093, chr14:95107608-95107761, chr14:95106041-95106223, chr14:95105678-95105783, chr14:95105071-95105246, chr14:95103346-95104126, chr14:95099780-95099935, chr14:95095825-95096713, chr14:95093888-95094156, chr14:95091203-95091365, chr14:95091034-95091109, chr14:95090496-95090663 |
| 28 | ***DROSHA*** | Full gene | NM_001382508.1 | ENST00000344624 | chr5:31529040-31529060, chr5:31526079-31526912, chr5:31521123-31521215, chr5:31515454-31515564, chr5:31514988-31515219, chr5:31511035-31511176, chr5:31508621-31508775, chr5:31504555-31504635, chr5:31495286-31495372, chr5:31493207-31493293, chr5:31486491-31486562, chr5:31484881-31484962, chr5:31483554-31483628, chr5:31472063-31472232, chr5:31467939-31468063, chr5:31466182-31466281, chr5:31464236-31464343, chr5:31451533-31451640, chr5:31449281-31449419, chr5:31448547-31448607, chr5:31437239-31437298, chr5:31435765-31435864, chr5:31431576-31431678, chr5:31429475-31429545, chr5:31424427-31424471, chr5:31422787-31422944, chr5:31421272-31421377, chr5:31410746-31410887, chr5:31409250-31409332, chr5:31409056-31409159, chr5:31406853-31406945, chr5:31405677-31405723, chr5:31401431-31401562 |
| 29 | ***DPYD*** | Hotspot | NM_000110.4 | ENST00000370192 | chr1:97450008-97450108, chr1:97579843-97579943, chr1:97573813-97573913, chr1:97515737-97515837, chr1:97082341-97082441 |
| 30 | ***EGFR*** | Hotspot | NM_005228.5 | ENST00000275493 | chr7:55143305-55143488, chr7:55154011-55154152, chr7:55160139-55160338, chr7:55165280-55165437, chr7:55173921-55174043, chr7:55174722-55174820, chr7:55181293-55181478, chr7:55191719-55191874 |
| 31 | ***EIF1AX*** | Full gene | NM_001412.4 | ENST00000379607 | chrX:20141625-20141642, chrX:20138539-20138622, chrX:20135738-20135841, chrX:20133957-20134007, chrX:20132182-20132263, chrX:20130516-20130607, chrX:20128304-20128311 |
| 32 | ***ELP1*** | Full gene | NM_003640.5 | ENST00000374647 | chr9:108930997-108931147, chr9:108929769-108929921, chr9:108927372-108927453, chr9:108926523-108926603, chr9:108922842-108922927, chr9:108919253-108919349, chr9:108918811-108918901, chr9:108917547-108917670, chr9:108916204-108916297, chr9:108912264-108912494, chr9:108911010-108911180, chr9:108908305-108908404, chr9:108906303-108906485, chr9:108903563-108903669, chr9:108902839-108902942, chr9:108901628-108901681, chr9:108901425-108901530, chr9:108900260-108900375, chr9:108899822-108899895, chr9:108898671-108898749, chr9:108898502-108898581, chr9:108897148-108897285, chr9:108896953-108897038, chr9:108896496-108896644, chr9:108893943-108894066, chr9:108892986-108893083, chr9:108891203-108891404, chr9:108889332-108889393, chr9:108882125-108882187, chr9:108881705-108881765, chr9:108880052-108880165, chr9:108879446-108879557, chr9:108878623-108878750, chr9:108877995-108878149, chr9:108874895-108874970, chr9:108869113-108869182 |
| 33 | ***ERBB2*** | Hotspot | NM_004448.4 | ENST00000269571 | chr17:39711928-39712047, chr17:39715446-39715536, chr17:39715740-39715939, chr17:39716301-39716433, chr17:39716515-39716605, chr17:39717320-39717480, chr17:39719787-39719834, chr17:39723319-39723457, chr17:39723538-39723660, chr17:39723912-39724010, chr17:39724726-39724911, chr17:39725049-39725204, chr17:39725327-39725402, chr17:39727689-39728046 |
| 34 | ***ERBB3*** | Full gene | NM_001982.4 | ENST00000267101 | chr12:56080301-56080382, chr12:56083751-56083902, chr12:56084995-56085181, chr12:56086531-56086656, chr12:56087577-56087642, chr12:56087795-56087913, chr12:56088021-56088162, chr12:56088543-56088656, chr12:56088748-56088868, chr12:56092747-56092820, chr12:56092986-56093076, chr12:56093345-56093550, chr12:56093764-56093896, chr12:56094099-56094189, chr12:56094402-56094556, chr12:56095257-56095310, chr12:56095665-56095806, chr12:56096503-56096622, chr12:56096748-56096846, chr12:56097045-56097230, chr12:56097785-56097940, chr12:56098500-56098575, chr12:56098759-56098905, chr12:56099648-56099745, chr12:56099838-56100029, chr12:56100174-56100245, chr12:56101061-56101361, chr12:56101529-56102055 |
| 35 | ***ESR1*** | Hotspot | NM_000125.4 | ENST00000206249 | chr6:151944173-151944508, chr6:152011656-152011794, chr6:152060991-152061124, chr6:152094385-152094568, chr6:152098732-152098966 |
| 36 | ***FBXW7*** | Full gene | NM_001349798.2 | ENST00000603548 | chr4:152411303-152411804, chr4:152350042-152350124, chr4:152346930-152347071, chr4:152337802-152337936, chr4:152332596-152332719, chr4:152330732-152330868, chr4:152329672-152329785, chr4:152328208-152328389, chr4:152326006-152326231, chr4:152324184-152324394, chr4:152322881-152323149 |
| 37 | ***FGFR1*** | Hotspot | NM_023110.3 | ENST00000447712 | chr8:38417306-38417416, chr8:38414779-38414901 |
| 38 | ***FGFR2*** | Hotspot | NM_000141.5 | ENST00000358487 | chr10:121519979-121520169, chr10:121515117-121515319, chr10:121498495-121498605, chr10:121487991-121488113 |
| 39 | ***FGFR3*** | Hotspot | NM_000142.5 | ENST00000440486 | chr4:1801835-1802025, chr4:1804330-1804520, chr4:1806051-1806173, chr4:1806546-1806683 |
| 40 | ***FGFR4*** | Hotspot | NM_213647.3 | ENST00000292408 | chr5:177089602-177089693, chr5:177090745-177090825, chr5:177093138-177093331, chr5:177095330-177095440, chr5:177095533-177095723 |
| 41 | ***FOXL2*** | Hotspot | NM_023067.4 | ENST00000648323 | chr3:138946275-138946374 |
| 42 | ***FRK*** | Hotspot | NM_002031.3 | ENST00000606080 | chr6:115944244-115944425 |
| 43 | ***GATA3*** | Full gene | NM_001002295.2 | ENST00000379328 | chr10:8055654-8055896, chr10:8058305-8058841, chr10:8063993-8064138, chr10:8069473-8069598, chr10:8073739-8074025 |
| 44 | ***GNA11*** | Hotspot | NM_002067.5 | ENST00000078429 | chr19:3114944-3115072, chr19:3118924-3119053 |
| 45 | ***GNAQ*** | Hotspot | NM_002072.5 | ENST00000286548 | chr9:77797520-77797648, chr9:77794463-77794592 |
| 46 | ***GNAS*** | Hotspot | NM_000516.7 | ENST00000371085 | chr20:58909162-58909216, chr20:58909350-58909423, chr20:58909521-58909579 |
| 47 | ***GPR161*** | Full gene | NM_001375883.1 | ENST00000682931 | chr1:168104477-168104850, chr1:168096508-168097232, chr1:168090564-168090668, chr1:168087585-168087704, chr1:168085529-168085796 |
| 48 | ***H3-3A*** | Hotspot | NM_002107.7 | ENST00000366815 | chr1:226064352-226064479 |
| 49 | ***H3-3B*** | Hotspot | NM_005324.5 | ENST00000254810 | chr17:75779047-75779174 |
| 50 | ***H3C2*** | Full gene | NM_003537.4 | ENST00000621411 | chr6:26031651-26032060 |
| 51 | ***H3C3*** | Full gene | NM_003531.3 | ENST00000612966 | chr6:26045411-26045821 |
| 52 | ***HNF1A*** | Full gene | NM_000545.8 | ENST00000257555 | chr12:120978769-120979094, chr12:120988833-120989032, chr12:120993520-120993706, chr12:120994164-120994405, chr12:120996262-120996413, chr12:120996541-120996742, chr12:120997474-120997665, chr12:120999268-120999389, chr12:120999483-120999627, chr12:121001065-121001194 |
| 53 | ***HRAS*** | Hotspot | NM_005343.4 | ENST00000311189 | chr11:534212-534322, chr11:533766-533944, chr11:533453-533612 |
| 54 | ***IDH1*** | Hotspot | NM_005896.4 | ENST00000345146 | chr2:208248369-208248660 |
| 55 | ***IDH2*** | Hotspot | NM_002168.4 | ENST00000330062 | chr15:90088587-90088747 |
| 56 | ***IL6ST*** | Hotspot | NM_002184.4 | ENST00000381298 | chr5:55964146-55964312, chr5:55956025-55956235 |
| 57 | ***JAK1*** | Hotspot | NM_002227.4 | ENST00000342505 | chr1:64845513-64845640, chr1:64844754-64844889 |
| 58 | ***JAK2*** | Hotspot | NM_004972.4 | ENST00000381652 | chr9:5069925-5070052, chr9:5073698-5073785 |
| 59 | ***KBTBD4*** | Hotspot | NM_018095.6 | ENST00000430070 | chr11:47572928-47573790 |
| 60 | ***KEAP1*** | Full gene | NM_203500.2 | ENST00000171111 | chr19:10499395-10500034, chr19:10491577-10492262, chr19:10489648-10489853, chr19:10489192-10489368, chr19:10486650-10486818 |
| 61 | ***KIT*** | Full gene | NM_000222.3 | ENST00000288135 | chr4:54658013-54658081, chr4:54695512-54695781, chr4:54698284-54698565, chr4:54699630-54699766, chr4:54703724-54703892, chr4:54707098-54707287, chr4:54709424-54709539, chr4:54723584-54723698, chr4:54725857-54726050, chr4:54727218-54727324, chr4:54727325-54727415, chr4:54727416-54727542, chr4:54727823-54727927, chr4:54728011-54728121, chr4:54729335-54729485, chr4:54731328-54731419, chr4:54731871-54731998, chr4:54733069-54733192, chr4:54736498-54736609, chr4:54736721-54736820, chr4:54737175-54737280, chr4:54738429-54738559 |
| 62 | ***KRAS*** | Hotspot | NM_004985.5 | ENST00000311936 | chr12:25245274-25245384, chr12:25227234-25227412, chr12:25225614-25225773 |
| 63 | ***MAP2K1*** | Full gene | NM_002755.4 | ENST00000307102 | chr15:66387348-66387427, chr15:66435027-66435237, chr15:66436746-66436892, chr15:66443280-66443357, chr15:66444656-66444707, chr15:66481755-66481879, chr15:66484990-66485191, chr15:66487228-66487292, chr15:66489215-66489276, chr15:66489718-66489763, chr15:66490502-66490615 |

| No. | Gene | Panel coverage | RefSeq transcript (NCBI) | Ensembl transcript ID | Target regions (GRCh38/hg38) |
|---|---|---|---|---|---|
| 64 | ***MDM2*** | Full gene | NM_002392.6 | ENST00000258149 | chr12:68808476-68808491, chr12:68809208-68809292, chr12:68813554-68813628, chr12:68816812-68816945, chr12:68820325-68820374, chr12:68824363-68824430, chr12:68824555-68824651, chr12:68828771-68828931, chr12:68835829-68835984, chr12:68836672-68836749, chr12:68839274-68839851 |
| 65 | ***MET*** | Hotspot | NM_001127500.3 | ENST00000318493 | chr7:116699085-116700284, chr7:116771498-116771654, chr7:116771655-116771848, chr7:116771849-116772139, chr7:116774881-116775111, chr7:116777389-116777469, chr7:116778776-116778957, chr7:116781988-116782097, chr7:116783304-116783469, chr7:116795655-116795791, chr7:116795887-116796124 |
| 66 | ***MRE11*** | Full gene | NM_005591.4 | ENST00000323929 | chr11:94492782-94492803, chr11:94490833-94490965, chr11:94485924-94486084, chr11:94479674-94479761, chr11:94478735-94478876, chr11:94476289-94476403, chr11:94471574-94471759, chr11:94470471-94470642, chr11:94467813-94467893, chr11:94464113-94464239, chr11:94460936-94461036, chr11:94459408-94459581, chr11:94456276-94456338, chr11:94447219-94447438, chr11:94445810-94445893, chr11:94437177-94437235, chr11:94435832-94435899, chr11:94429911-94429986, chr11:94420124-94420181 |
| 67 | ***MTAP*** | Full gene | NM_002451.4 | ENST00000644715 | chr9:21802749-21802781, chr9:21815433-21815519, chr9:21816714-21816772, chr9:21818035-21818202, chr9:21837908-21838010, chr9:21854631-21854870, chr9:21859303-21859425, chr9:21861976-21862016 |
| 68 | ***MYC*** | Full gene | NM_002467.6 | ENST00000621592 | chr8:127736592-127736623, chr8:127738248-127739019, chr8:127740396-127740960 |
| 69 | ***MYCN*** | Full gene | NM_005378.6 | ENST00000281043 | chr2:15942064-15942854, chr2:15945493-15946099 |
| 70 | ***MYOD1*** | Full gene | NM_002478.5 | ENST00000250003 | chr11:17719782-17720412, chr11:17720902-17720980, chr11:17721255-17721510 |
| 71 | ***NBN*** | Full gene | NM_002485.5 | ENST00000265433 | chr8:89984525-89984563, chr8:89982722-89982855, chr8:89981375-89981523, chr8:89980734-89980893, chr8:89978220-89978323, chr8:89971173-89971290, chr8:89970364-89970557, chr8:89964410-89964507, chr8:89958725-89958854, chr8:89955283-89955555, chr8:89953244-89953691, chr8:89947824-89947892, chr8:89946140-89946295, chr8:89943253-89943366, chr8:89937026-89937075, chr8:89935580-89935612 |
| 72 | ***NF1*** | Full gene | NM_001042492.3 | ENST00000358273 | chr17:31095308-31095369, chr17:31155983-31156126, chr17:31159010-31159093, chr17:31163186-31163376, chr17:31169891-31169997, chr17:31181422-31181489, chr17:31181710-31181785, chr17:31182508-31182665, chr17:31200422-31200595, chr17:31201037-31201159, chr17:31201411-31201485, chr17:31206240-31206371, chr17:31214451-31214585, chr17:31219005-31219118, chr17:31221850-31221929, chr17:31223444-31223567, chr17:31225095-31225250, chr17:31226435-31226684, chr17:31227218-31227291, chr17:31227523-31227606, chr17:31229025-31229465, chr17:31229835-31229974, chr17:31230260-31230382, chr17:31230842-31230925, chr17:31232073-31232189, chr17:31232700-31232881, chr17:31233002-31233213, chr17:31235611-31235772, chr17:31235918-31236021, chr17:31248984-31249119, chr17:31252938-31253000, chr17:31258344-31258502, chr17:31259032-31259129, chr17:31260369-31260515 |
| 73 | ***NF2*** | Full gene | NM_000268.4 | ENST00000338641 | chr22:29603997-29604112, chr22:29636751-29636876, chr22:29639090-29639212, chr22:29642202-29642285, chr22:29654657-29654725, chr22:29655594-29655676, chr22:29658189-29658264, chr22:29661205-29661339, chr22:29664990-29665064, chr22:29668333-29668446, chr22:29671826-29671948, chr22:29673269-29673486, chr22:29674836-29674941, chr22:29678196-29678323, chr22:29681439-29681601, chr22:29694752-29694803 |
| 74 | ***NOTCH1*** | Full gene | NM_017617.5 | ENST00000651671 | chr9:136545726-136545788, chr9:136544024-136544102, chr9:136523717-136523979, chr9:136522850-136523188, chr9:136519443-136519565, chr9:136518591-136518824, chr9:136518137-136518292, chr9:136517752-136517937, chr9:136517272-136517385, chr9:136515981-136516094, chr9:136515483-136515716, chr9:136515290-136515400, chr9:136514510-136514702, chr9:136513392-136513537, chr9:136513021-136513134, chr9:136511152-136511271, chr9:136510653-136510805, chr9:136509733-136509961, chr9:136508870-136509071, chr9:136508232-136508385, chr9:136507955-136508139, chr9:136507305-136507437, chr9:136506716-136506973, chr9:136506527-136506639, chr9:136505310-136505881, chr9:136504673-136505104, chr9:136503182-136503330, chr9:136502272-136502488, chr9:136502001-136502088, chr9:136501748-136501913, chr9:136500552-136500847, chr9:136499112-136499259, chr9:136498899-136498996, chr9:136496099-136497558 |
| 75 | ***NRAS*** | Hotspot | NM_002524.5 | ENST00000369535 | chr1:114716050-114716160, chr1:114713800-114713978, chr1:114709569-114709728 |
| 76 | ***NTRK1*** | Hotspot | NM_002529.4 | ENST00000524377 | chr1:156876400-156876572, chr1:156879122-156879362 |
| 77 | ***NTRK2*** | Hotspot | NM_001018064.3 | ENST00000376213 | chr9:84948462-84948634, chr9:84955283-84955517 |
| 78 | ***NTRK3*** | Hotspot | NM_001012338.3 | ENST00000360948 | chr15:87933012-87933184, chr15:87929191-87929434 |
| 79 | ***PALB2*** | Full gene | NM_024675.4 | ENST00000261584 | chr16:23641110-23641160, chr16:23638070-23638129, chr16:23637850-23637952, chr16:23634862-23636334, chr16:23629640-23630469, chr16:23629204-23629275, chr16:23626236-23626397, chr16:23624009-23624094, chr16:23622969-23623130, chr16:23621362-23621478, chr16:23614004-23614091, chr16:23607864-23608012, chr16:23603457-23603669 |
| 80 | ***PDGFRA*** | Hotspot | NM_006206.6 | ENST00000257290 | chr4:54274841-54274973, chr4:54277388-54277492, chr4:54277896-54278006, chr4:54278362-54278515, chr4:54285841-54285963 |
| 81 | ***PDGFRB*** | Hotspot | NM_002609.4 | ENST00000261799 | chr5:150126520-150126614, chr5:150125445-150125577, chr5:150124727-150124831, chr5:150124250-150124360, chr5:150123042-150123201, chr5:150121880-150122040, chr5:150121204-150121322, chr5:150120888-150121010, chr5:150120012-150120123, chr5:150119467-150119566, chr5:150118747-150118852 |
| 82 | ***PIK3CA*** | Full gene | NM_006218.4 | ENST00000263967 | chr3:179198827-179199177, chr3:179199690-179199899, chr3:179201290-179201540, chr3:179203544-179203789, chr3:179204503-179204588, chr3:179209595-179209700, chr3:179210186-179210338, chr3:179210431-179210565, chr3:179218210-179218334, chr3:179219196-179219277, chr3:179219571-179219735, chr3:179219949-179220052, chr3:179220986-179221157, chr3:179224081-179224187, chr3:179224700-179224821, chr3:179225962-179226040, chr3:179229272-179229442, chr3:179230004-179230121, chr3:179230225-179230376, chr3:179234094-179234364 |
| 83 | ***PIK3R1*** | Full gene | NM_181523.3 | ENST00000521381 | chr5:68226676-68227009, chr5:68273390-68273482, chr5:68273939-68274013, chr5:68279602-68279733, chr5:68280528-68280729, chr5:68280927-68281006, chr5:68292259-68292361, chr5:68293101-68293199, chr5:68293303-68293483, chr5:68293709-68293834, chr5:68294536-68294678, chr5:68295148-68295324, chr5:68295420-68295488, chr5:68296171-68296341, chr5:68297412-68297603 |
| 84 | ***POLE*** | Hotspot | NM_006231.4 | ENST00000320574 | chr12:132676546-132676653, chr12:132676094-132676204, chr12:132675735-132675820, chr12:132675398-132675517, chr12:132673575-132673707, chr12:132673164-132673277 |
| 85 | ***PRKCA*** | Hotspot | NM_002737.3 | ENST00000413366 | chr17:66742622-66742760 |
| 86 | ***PTCH1*** | Full gene | NM_000264.5 | ENST00000331920 | chr9:95508161-95508363, chr9:95506407-95506599, chr9:95485685-95485874, chr9:95482134-95482203, chr9:95481949-95482040, chr9:95480390-95480588, chr9:95479969-95480090, chr9:95479000-95479147, chr9:95478055-95478186, chr9:95477547-95477702, chr9:95476759-95476857, chr9:95476034-95476159, chr9:95469813-95469931, chr9:95468751-95469153, chr9:95467116-95467425, chr9:95461856-95461998, chr9:95459600-95459783, chr9:95458013-95458293, chr9:95456276-95456413, chr9:95453478-95453620, chr9:95449841-95449940, chr9:95449069-95449323, chr9:95446911-95447451 |
| 87 | ***PTEN*** | Full gene | NM_000314.8 | ENST00000371953 | chr10:87864470-87864548, chr10:87894025-87894109, chr10:87925513-87925557, chr10:87931046-87931089, chr10:87933013-87933251, chr10:87952118-87952259, chr10:87957853-87958019, chr10:87960894-87961118, chr10:87965287-87965472 |
| 88 | ***PTPN11*** | Full gene | NM_002834.5 | ENST00000351677 | chr12:112419110-112419125, chr12:112446276-112446398, chr12:112450318-112450512, chr12:112453195-112453387, chr12:112454564-112454680, chr12:112455950-112456063, chr12:112472944-112473040, chr12:112477651-112477730, chr12:112477857-112478015, chr12:112482074-112482205, chr12:112486475-112486629, chr12:112488443-112488510, chr12:112489024-112489175, chr12:112502144-112502256, chr12:112504695-112504766 |
| 89 | ***RAD50*** | Full gene | NM_005732.4 | ENST00000378823 | chr5:132557323-132557453, chr5:132559284-132559367, chr5:132575777-132575928, chr5:132579317-132579502, chr5:132579862-132580066, chr5:132587562-132587690, chr5:132587924-132588089, chr5:132588687-132588880, chr5:132589631-132589837, chr5:132591224-132591406, chr5:132591877-132592034, chr5:132594869-132595044, chr5:132595573-132595810, chr5:132603300-132603489, chr5:132603920-132604046, chr5:132604806-132604999, chr5:132608615-132608725, chr5:132609117-132609209, chr5:132609283-132609396, chr5:132616003-132616130, chr5:132618070-132618294, chr5:132637115-132637200, chr5:132638081-132638223, chr5:132640672-132640805 |
| 90 | ***RAD51B*** | Full gene | NM_133510.4 | ENST00000471583 | chr14:67823544-67823627, chr14:67825464-67825577, chr14:67835080-67835196, chr14:67865003-67865139, chr14:67885869-67885988, chr14:67887021-67887204, chr14:68291884-68291980, chr14:68411424-68411527, chr14:68468172-68468250, chr14:68477648-68477666 |
| 91 | ***RAD51C*** | Full gene | NM_058216.3 | ENST00000337432 | chr17:58692643-58692788, chr17:58694931-58695189, chr17:58696693-58696859, chr17:58703196-58703329, chr17:58709859-58709990, chr17:58720746-58720812, chr17:58724040-58724100, chr17:58732484-58732544, chr17:58734118-58734224 |
| 92 | ***RAD51D*** | Full gene | NM_002878.4 | ENST00000345365 | chr17:35119532-35119614, chr17:35119111-35119172, chr17:35118501-35118619, chr17:35107366-35107447, chr17:35106988-35107122, chr17:35106386-35106481, chr17:35103454-35103544, chr17:35103254-35103324, chr17:35101201-35101365, chr17:35100951-35101036 |
| 93 | ***RB1*** | Full gene | NM_000321.3 | ENST00000267163 | chr13:48303913-48304049, chr13:48307280-48307406, chr13:48342599-48342714, chr13:48345080-48345199, chr13:48347825-48347863, chr13:48348956-48349023, chr13:48360017-48360127, chr13:48362815-48362957, chr13:48364894-48364971, chr13:48367494-48367603, chr13:48368527-48368604, chr13:48373405-48373492, chr13:48376918-48377034, chr13:48379594-48379650, chr13:48380053-48380084, chr13:48380165-48380241, chr13:48381247-48381443, chr13:48452993-48453111, chr13:48456204-48456349, chr13:48459688-48459833, chr13:48463731-48463835, chr13:48464998-48465111, chr13:48465205-48465368, chr13:48473360-48473390, chr13:48476701-48476843, chr13:48477355-48477404, chr13:48479998-48480071 |
| 94 | ***RET*** | Full gene | NM_020975.6 | ENST00000355710 | chr10:43077257-43077331, chr10:43100459-43100722, chr10:43102342-43102629, chr10:43104952-43105193, chr10:43106376-43106571, chr10:43109031-43109230, chr10:43111207-43111465, chr10:43112099-43112224, chr10:43112583-43112963, chr10:43113556-43113675, chr10:43114480-43114736, chr10:43116584-43116731, chr10:43118373-43118480, chr10:43119531-43119745, chr10:43120081-43120203, chr10:43121946-43122016, chr10:43123671-43123808, chr10:43124883-43124982 |

| No. | Gene | Panel coverage | RefSeq transcript (NCBI) | Ensembl transcript ID | Target regions (GRCh38/hg38) |
|---|---|---|---|---|---|
| 95 | ***RNF43*** | Full gene | NM_017763.6 | ENST00000407977 | chr17:58415326-58415577, chr17:58370911-58371033, chr17:58363526-58363600, chr17:58363275-58363406, chr17:58362544-58362648, chr17:58360783-58360944, chr17:58360149-58360251, chr17:58357468-58358823, chr17:58354943-58354986 |
| 96 | ***ROS1*** | Hotspot | NM_002944.3 | ENST00000368508 | chr6:117319868-117320030, chr6:117318188-117318252, chr6:117317143-117317272, chr6:117308794-117308928 |
| 97 | ***SDHA*** | Full gene | NM_004168.4 | ENST00000264932 | chr5:218355-218418, chr5:223482-223568, chr5:224360-224521, chr5:225419-225562, chr5:225883-226047, chr5:228185-228333, chr5:230876-231000, chr5:233477-233645, chr5:235144-235339, chr5:236428-236599, chr5:240358-240476, chr5:250992-251103, chr5:251338-251468, chr5:254393-254506, chr5:256334-256422 |
| 98 | ***SDHB*** | Full gene | NM_003000.3 | ENST00000375499 | chr1:17053948-17054018, chr1:17044761-17044888, chr1:17033060-17033145, chr1:17028600-17028736, chr1:17027749-17027865, chr1:17023973-17024074, chr1:17022608-17022730, chr1:17018880-17018958 |
| 99 | ***SDHC*** | Full gene | NM_003001.5 | ENST00000367975 | chr1:161314405-161314425, chr1:161323614-161323670, chr1:161328396-161328497, chr1:161340594-161340655, chr1:161356677-161356840, chr1:161362329-161362435 |
| 100 | ***SDHD*** | Full gene | NM_003002.4 | ENST00000375549 | chr11:112086907-112086959, chr11:112087857-112087973, chr11:112088867-112089011, chr11:112094805-112094972 |
| 101 | ***SF3B1*** | Hotspot | NM_012433.4 | ENST00000335508 | chr2:197402554-197402828, chr2:197401983-197402132, chr2:197401740-197401890, chr2:197400713-197400938 |
| 102 | ***SMAD4*** | Full gene | NM_005359.6 | ENST00000342988 | chr18:51047047-51047295, chr18:51048686-51048860, chr18:51049295-51049324, chr18:51054781-51054993, chr18:51058125-51058244, chr18:51058340-51058456, chr18:51059866-51059916, chr18:51065423-51065606, chr18:51067019-51067187, chr18:51076638-51076776, chr18:51078256-51078467 |
| 103 | ***SMARCA4*** | Full gene | NM_001128844.3 | ENST00000429416 | chr19:10984151-10984373, chr19:10985273-10985405, chr19:10986189-10986593, chr19:10986905-10987003, chr19:10987666-10987924, chr19:10989317-10989443, chr19:10991150-10991323, chr19:10994828-10995001, chr19:10996213-10996380, chr19:10996494-10996544, chr19:11003029-11003159, chr19:11003340-11003397, chr19:11007902-11008023, chr19:11010381-11010531, chr19:11012949-11013112, chr19:11018957-11019023, chr19:11019591-11019701, chr19:11021725-11021967, chr19:11023518-11023631, chr19:11024331-11024438, chr19:11025422-11025508, chr19:11026300-11026346, chr19:11027784-11027950, chr19:11030730-11030893, chr19:11033290-11033517, chr19:11033767-11033865, chr19:11034123-11034200, chr19:11034914-11035132, chr19:11041307-11041560, chr19:11058255-11058363, chr19:11058788-11058889, chr19:11059753-11059885, chr19:11060045-11060187, chr19:11061784-11061818 |
| 104 | ***SMARCB1*** | Full gene | NM_003073.5 | ENST00000644036 | chr22:23787168-23787262, chr22:23791756-23791894, chr22:23793559-23793688, chr22:23800944-23801081, chr22:23803295-23803422, chr22:23816770-23816936, chr22:23825225-23825415, chr22:23833572-23833703, chr22:23834141-23834182 |
| 105 | ***SMO*** | Full gene | NM_005631.5 | ENST00000249373 | chr7:129189152-129189482, chr7:129203384-129203589, chr7:129205203-129205412, chr7:129205610-129205782, chr7:129206150-129206369, chr7:129206464-129206587, chr7:129208759-129208851, chr7:129209289-129209397, chr7:129210363-129210548, chr7:129210965-129211113, chr7:129211636-129211770, chr7:129212024-129212453 |
| 106 | ***SPOP*** | Hotspot | NM_003563.3 | ENST00000393328 | chr17:49619234-49619385, chr17:49618981-49619108 |
| 107 | ***STAT3*** | Hotspot | NM_139276.3 | ENST00000264657 | chr17:42346569-42346713, chr17:42338731-42338812, chr17:42324711-42324846, chr17:42323004-42323143, chr17:42322282-42322494 |
| 108 | ***STK11*** | Full gene | NM_000455.5 | ENST00000326873 | chr19:1206914-1207203, chr19:1218417-1218500, chr19:1219324-1219413, chr19:1220373-1220505, chr19:1220581-1220717, chr19:1221213-1221340, chr19:1221949-1222006, chr19:1222985-1223172, chr19:1226454-1226647 |
| 109 | ***SUFU*** | Full gene | NM_016169.4 | ENST00000369902 | chr10:102504151-102504334, chr10:102509169-102509303, chr10:102549970-102550106, chr10:102592582-102592724, chr10:102593636-102593721, chr10:102593993-102594065, chr10:102597140-102597293, chr10:102599433-102599544, chr10:102615268-102615402, chr10:102617290-102617428, chr10:102627175-102627243, chr10:102630066-102630157 |
| 110 | ***TERT*** | Hotspot | NM_198253.3 | ENST00000310581 | chr5:1295016-1295205 |
| 111 | ***TP53*** | Full gene | NM_000546.6 | ENST00000269305 | chr17:7676521-7676594, chr17:7676382-7676403, chr17:7675994-7676272, chr17:7675053-7675236, chr17:7674859-7674971, chr17:7674181-7674290, chr17:7673701-7673837, chr17:7673535-7673608, chr17:7670609-7670715, chr17:7669609-7669690, chr17:7673219-7673339 |
| 112 | ***VHL*** | Full gene | NM_000551.4 | ENST00000256474 | chr3:10141848-10142187, chr3:10146514-10146636, chr3:10149787-10149965 |
